# IPF distal lung epithelial cells acquire a DNA methylation signature consistent with activation of basal cell transcriptional programs

**DOI:** 10.64898/2026.09.18.752396

**Authors:** Arlo C. Colvard, Stefano Iantorno, Isabella P. Gaona, Merced Malabanan, Taylor Sherrill, Rafael J. Fernandez, Ciara M. Shaver, Lorraine B. Ware, Bradley W. Richmond, Jennifer MS Sucre, Nicholas E. Banovich, A. Scott McCall, Jonathan A. Kropski, Jason J. Gokey

## Abstract

Idiopathic pulmonary fibrosis (IPF) is a progressive fibrotic interstitial lung disease associated with failed alveolar epithelial repair with an expansion of aberrant airway-like epithelium in the alveolar space leading to lung function decline usually resulting in death within 3-5 years of diagnosis. While single-cell and spatial transcriptomic approaches have been used to characterize disease-emergent cell populations, less is known about the regulation of transcriptional programs that drive failed alveolar epithelial cell repair in IPF. DNA methylation is a fundamental layer of gene regulation that stabilizes differentiated cell identity; however, changes in methylation in the epithelial compartment in IPF have not been studied. To identify novel epigenetic mediators of epithelial cell dysfunction, we performed high-resolution DNA methylation profiling of purified distal lung epithelial cells from 10 age-matched control and 14 IPF lungs using Oxford Nanopore Technologies (ONT) whole-genome, long-read sequencing. We identified widespread methylome remodeling in the IPF lung epithelium, with 84% of differentially methylated regions (DMRs) hypomethylated. DMRs were largely found outside of promoters, with 88% outside of ±3 kb from the transcription start site (TSS), consistent with altered distal regulatory element activity. DMRs were enriched for transcription factor (TF) binding site motifs and gene associations consistent with dysregulation of polycomb repressive complex 2 (PRC2) and increases in p63 activity. DNA methylation at DMRs associated with p63 target genes, including *KRT5*, and genes implicated in failed epithelial repair, including *MUC5B, MMP7, and S100A2* inversely correlated with gene expression. Consistent with the observed dysregulation of the IPF epithelial methylome, an experimental co-culture model of alveolar type II epithelial cell (AT2s) to basal-like epithelial cell transdifferentiation revealed widespread hypomethylation. Further, sites associated with NKX2.1 and FOXA1/2 binding, TFs involved in alveolar fate maintenance, were hypermethylated, suggesting loss of epigenetic regulation of alveolar identity. Together, these data implicate DNA methylation in the failed alveolar epithelial repair processes in IPF, potentially providing future therapeutic strategies by identifying putative regulatory elements associated with aberrant transcriptional programs.

## Introduction

Idiopathic pulmonary fibrosis (IPF) is a progressive interstitial lung disease characterized by extensive fibrotic remodeling of the lung parenchyma, loss of functional alveolar epithelium, and a progressive decline in respiratory function^1^. IPF is thought to result from recurrent lung epithelial injury followed by aberrant repair and persistent profibrotic signaling, resulting in excessive extracellular matrix deposition. The cellular and molecular mechanisms that drive failed repair remain incompletely understood. Current FDA-approved therapies for IPF only slow disease progression, with lung transplantation remaining the sole management option^1–3^. Therefore, an improved understanding of the regulatory mechanisms underlying failed epithelial differentiation and repair, including dysregulation of upstream transcription factor activity, in IPF is urgently needed and may inform the development of targeted therapies.

Previous studies have employed a range of multi-omics techniques to elucidate pathways and mediators associated with disease progression. Single-cell RNA sequencing studies have revealed the emergence of conducting airway populations, including goblet, secretory, ciliated, basal, and KRT5-/KRT17+ cells in the distal lung in IPF ^4–6^. *p63*+ basal cells are the primary stem cells of the airway and, under homeostatic conditions, serve to maintain epithelial populations following injury. However, the expansion of *p63*+ cells in IPF distal lung and their acquisition of a *KRT17*-high and *PTEN-*low phenotype are concomitant with a loss of functional alveolar epithelium, activation of fibroblast populations and the deposition of excessive amounts of fibrotic extracellular matrix^7^. IPF *KRT5*+ basal cells are also transcriptionally distinct from those found in control lungs, enhance ECM production by fibroblasts in co-culture models, and cause lung fibrosis in mice^7^. Together, basal cells and other airway populations replace the functional alveolar epithelium, leading to reduced respiratory function. While the origin of these cells remains to be clearly defined, recent research has shown that *p63*+ basal and KRT5-/KRT17+ cell populations can emerge from transdifferentiation of type-2 alveolar epithelial cells (AT2s) following co-culture with lung mesenchyme, suggesting a similar process may drive the disease-associated shifts in the epithelial repertoire of the alveoli^8^. Single-cell transcriptomic studies have allowed for characterization of the expression profiles of ectopic, disease-emergent cell populations in IPF; however, the regulatory mechanisms that permit and underpin this aberrant epithelial differentiation, predisposing the lung to failed repair, remain poorly understood.

DNA methylation is a fundamental layer of gene regulation, and the elucidation of its precise alterations in IPF may allow for the development of therapies that target mediators of the maladaptive transcriptional reprogramming in the lung epithelium. Previous studies of the methylome of the IPF lung have relied on array-based profiling of bulk lung tissue, an approach that presents numerous challenges for accurate characterization of the regulatory role for DNA methylation in IPF, including confounding factors such as cell type heterogeneity, biased CpG selection, and a focus on individual CpGs rather than broader regulatory regions^9^. As genome-wide methylation profiling studies integrated with transcriptomic analyses have revealed a regulatory role for cell-type specific methylation at intragenic regions, distal regulatory elements, and transcription factor binding sites, similar analytical techniques are necessary to comprehensively characterize the role of methylation in regulating aberrant epithelial differentiation in IPF^10–18^. Here, we used Oxford Nanopore Technologies (ONT) long-read sequencing to perform high-resolution characterization of CpG methylation and describe region-based changes in methylation states in isolated lung epithelial cells from IPF subjects and age-matched control donors.

We hypothesized that alterations in DNA methylation at regulatory regions in the distal lung epithelium allow for activation of aberrant transcriptional programs associated with loss of lineage fidelity. To investigate this, we leveraged both a sliding window analysis of methylation at promoters and a segmentation approach designed to capture stable patterns of methylation across consecutive CpGs at any position in the genome.

Our data reveal changes in DNA methylation in putative regulatory regions accompanying activation of basal cell transcriptional programs in the IPF distal lung epithelium. We identify the location of putative regulatory elements that are differentially methylated with failed alveolar epithelial repair, including elements overlapping IPF risk variants, and suggest potential regulatory relationships with differentially expressed genes. We further describe TFs predicted to act in these regions of altered methylation that may serve as targetable mediators of aberrant epithelial differentiation.

## Methods

### Subjects and samples

Peripheral lung tissue used for CD326+ cell selection was obtained from control donor lungs (5 male, 5 female) deemed unsuitable for transplantation and from subjects with idiopathic pulmonary fibrosis (IPF) who underwent lung transplantation (7 male, 7 female) at Vanderbilt University Medical Center. Lung epithelium was isolated fresh (within 48 hours of procurement) from peripheral lung regions or from minced peripheral lung tissue cryopreserved in 90% fetal bovine serum and 10% dimethyl sulfoxide. No difference in DNA methylation was observed in frozen relative to fresh samples. This study was approved by the Institutional Review Board at Vanderbilt University (#060165, #171657, #192004).

Peripheral lung tissue used for HTII-280+ cell selection was obtained from donor lungs that were rejected for transplantation at UCSF. Tissue from peripheral lung regions was processed fresh.

### CD326+ cell selection and DNA and RNA isolation

Minced tissue was homogenized in DMEM (Gibco, #31053028) containing collagenase I (125 CDU/mL, Sigma-Aldrich, C0130), dispase II (1U/mL, Roche, 04942078001), DNase I (2,500 DU/mL, Millipore Sigma, 260913-10MU), leupeptin (50 nM, RPI, L22035) solution for 15 minutes in a gentleMACs Octo Dissociator with constant stirring at a temperature of 37 C. The resulting suspension was then passed through sterile gauze, 100 µm, and 70µm filters (MTC Bio, C4100 and C4070) to generate single-cell suspensions. CD326+ epithelial cells were enriched using LS Column (Miltenyi Biotec, 130-042-401) positive selection with CD326 MicroBeads (Miltenyi Biotec, 130-061-101) according to the manufacturer’s protocol. DNA was isolated from CD326+ epithelium according to the DNeasy Blood & Tissue Kit (QIAGEN, 69504) instructions. RNA was isolated from CD326+ epithelium according to the RNeasy Plus Mini Kit (QIAGEN, 74136).

### HTII-280 fluorescence activated sorting

Lung digestion and fluorescence activated sorting for HTII-280+ cells was performed as previously described^8^. Briefly, human lung pieces were washed in PBS and HBSS, minced into 1 cm^3^ pieces, digested for 2h at 37 ℃ in a HBSS solution containing Dispase II (15 U/mL), collagenase type I (225 U/mL), Dnase I (100 U/mL), and 1% Pen/Strep, with (1:400) Fungizone added for the final 30 minutes. Single cell suspensions were generated from digested tissue in a blender, filtered through sterile gauze, 100 µm, 70 µm, and 40 µm filters, and treated with a red blood cell lysis buffer. FcR blocking was performed and immune and endothelial cells were depleted using CD45, CD31, and CD11b biotinylated antibodies and streptavidin beads (25 µL/mL). AT2s were then sorted as live/EpCAM+/HTII-280+ cells, and adult human lung mesenchyme (AHLM) cells were sorted as live/CD45−/CD11b−/CD31−/EpCAM− cells.

### Cell culture and DNA isolation

AT2s and AHLM cells were co-cultured as previously described^8^. Briefly, AT2 organoids and mesenchymal cells were co-cultured at a ratio of 5,000:30,000 in modified MTEC media and growth factor-reduced Matrigel at a 1:1 ratio. The cell-Matrigel mixture was plated on a transwell and incubated with 10 µM ROCK inhibitor for 48 hours. Following 14 days of co-culture, the cell-Matrigel mixture was washed with PBS and incubated in the lung digestion cocktail used for HTII-280 fluorescence activated sorting for 1h at 37 ℃ with periodic resuspension. The resulting suspension was resuspended in TrypLE and shaken at 37 ℃ for 20 minutes. FcR blocking was performed and epithelium was isolated using CD326 biotinylated antibodies and streptavidin beads (1:50). DNA was isolated from CD326+ epithelium according to the DNeasy Blood & Tissue Kit (QIAGEN, 69504) instructions.

### Long-read sequencing and basecalling

Genomic DNA was prepared for long-read sequencing according to the Ligation sequencing DNA V14 (SQK-LSK114) protocol from Oxford Nanopore Technologies. Samples were loaded onto FLO-PRO114M PromethION flow cells and sequenced on a PromethION 2 Solo. An average genome-wide coverage of 30x was targeted. Following the completion of sequencing, flow cells with a pore count > 2000 were washed and reloaded with DNA from an additional subject according to the Flow Cell Wash Kit (EXP-WSH004) instructions. The raw sequencing POD5 files from the CD326+ isolation from IPF and control subjects were basecalled using *Dorado* with the dna_r10.4.1_e8.2_400bps_sup@v5.0.0 model. The raw sequencing POD5 files from the co-culture model samples were basecalled using *Dorado* with the recently released dna_r10.4.1_e8.2_400bps_sup@v5.2.0 model.

### Preprocessing and quality control

Alignment statistics for each subject were obtained from the alignment report generated through the *epi2me* Human Variation workflow. Bedmethyl files were generated using pileup from the *modkit* version 0.6.1 package with the following options: modifications were filtered to include the 5mC modification (--modified-bases 5mC), modifications were aggregated across CpGs on the + and - strands (--combine strands), and modifications were restricted to those found at CpG dinucleotides (--cpg). The 10th percentile of the overall distribution of confidence scores for base identities was used as a minimum threshold for each subject independently. For inclusion in analysis, CpGs were filtered for ≥ 5x coverage in 90% of samples in each group (≥ 5x coverage in ≥ 9 control and ≥ 12 IPF subjects).

### Genomic region methylation analysis and annotation

Promoters were defined using the *TxDb.Hsapiens.UCSC.hg19.knownGene* database as 3,000 bp surrounding each transcription start site. Promoter regions were subdivided into twelve consecutive 500-bp tiles and used to annotate CpG level methylation counts. Differential methylation was tested through *methylKit*^19^ using a Chi-squared test on a logistic regression model of methylation with a multinomial overdispersion correction and Benjamini-Hochberg correction for multiple comparisons. Sex and reported smoking history, defined as ever or never smokers, were included as covariates. We defined DMR significance as FDR ≤ 0.01 and absolute methylation difference ≥ 10%, consistent with previously described biologically relevant alterations^20^.

Genome-wide unmethylated regions (UMRs), low-methylated regions (LMRs), and differentially methylated regions (DMRs) were identified used *MethyLasso*^21^ with the following options: a replicate coverage score of 0.9 was required for a region to be included (-r 0.9) and all regions tested for differential methylation were included in the output DMR file (-p 1 and -d 0). UMRs and LMRs present in control and IPF groups were identified separately. Statistical analysis was conducted on all regions tested for differential methylation by *MethyLasso* through *methylKit*.

The region annotations of control and IPF LMRs and UMRs from *MethyLasso* were merged using *GenomicRanges::reduce*. The resulting LMR and UMR regions were used to annotate CpG level methylation counts in *methylKit*. DMR annotations were separately used to annotate methylation counts in *methylKit*. Regions were assigned to the nearest-neighbor TSS using the *ChIPseekr* and *TxDb.Hsapiens.UCSC.hg19.knownGene* R packages.

### Transcription factor binding site motif (TFBS) and gene set enrichment analysis

Using *homer2*, we identified TFBS motif enrichment at DMRs with increased or decreased methylation in IPF, separately. This tool identifies motifs overrepresented in genomic intervals of interest by comparison with a background set of intervals with similar nucleotide and CpG composition. P-value was adjusted for multiple comparisons using the Benjamini-Hochberg correction. Enriched TFs were plotted according to the -log_10_ transformation of the FDR-adjusted p-value and the quotient of the % of target sequences containing motif and % of background sequences containing motifs.

Using *enrichR*^22–24^, we evaluated TF binding site enrichment of genes associated with DMRs with increased or decreased methylation in IPF separately. Gene lists were compared to the *ENCODE_and_ChEA_Consensus_TFs_from_ChIP-X* library to perform gene set enrichment. The -log_10_ of FDR-adjusted p-values were plotted against the odds ratio for each TF.

### RNA-sequencing analysis

When available, an extra sample was collected and RNA was isolated from CD326+ cells isolated from control (male n=3, female n=3) and IPF (male n=6, female n=5) subject lungs according to the Qiagen RNeasy Plus Mini Kit protocol. RNA sequencing was performed by Azenta Life Sciences targeting 30M paired-end reads. *Trimmomatic* version 0.39 was used to remove adapter sequences. *STAR*^25^ version 2.7.11b was used to align reads to the GENCODE GRCh38 human genome assembly. *Subread*^26^ version 2.1.1 was used to obtain feature counts by gene. Genes were included in downstream analysis if expression was detected in ≥ 5 control or ≥ 8 IPF subjects. *DESeq2*^27^ was used to identify statistically significant changes in expression between control and IPF subjects, normalize counts, and perform variance-stabilized transformation. Differentially expressed genes were defined as log_2_(fold change) ≥ 1 and an FDR-adjust p-value ≤ 0.05.

### eQTM analysis

Expression quantitative trait methylation (eQTMs) were identified using the R package *Matrix eQTL*^28^. Methylation percentages at DMRs were substituted for SNPs in the pipeline for this package. DMR location was defined as the midpoint of each region. The variance-stabilized, normalized counts of differentially expressed genes (FDR ≤ 0.05 and log_2_(fold-change) ≥ 1) from RNA-sequencing were used as the input for the expression matrix. Reported smoking history and subject sex were included as covariates for this analysis. For eQTM analysis of DMRs obtained from *methyLasso*, a DMR to gene distance maximum threshold of 1 million base pairs was set. The TSS was standardized as the gene start for each gene.

### DMR-WAS

We obtained IPF GWAS summary statistics from Chin et al. 2026 (https://github.com/genomicsITER/PFgenetics; doi: 10.1183/13993003.00506-2026)29 and identified direct overlaps of variants with DMRs using *GenomicRanges*. We used *rtracklayer::liftOver* to transpose DMRs from the hg19 to the hg38 genome build. For both overlapping and nonoverlapping SNPs, the total number of variants (25,552,573) were used for a Bonferroni correction of a significance threshold of 0.05, giving a genome-wide significance threshold of 1.96^-9^.

### Statistical analysis

To account for sex as a biological variable, we used equal numbers of male and female subjects for DNA methylation analysis and approximately equal numbers for RNA-seq analysis.

For both our promoter-restricted and genome-wide DMR analysis, we used *methylKit* with Chi-squared testing, multinomial overdispersion correction, Benjamini-Hochberg multiple testing correction, and sex and reported smoking history as covariates. We defined DMR significance as FDR ≤ 0.01 and absolute methylation difference ≥ 10%.

We assessed differential gene expression using DESeq2, with DEGs defined as FDR ≤ 0.05 and absolute log_2_(fold change) ≥ 1 and sex and reported smoking history as covariates. We used *Matrix eQTL* to identify eQTMs with sex and reported smoking history as covariates; we defined significant associations as FDR ≤ 0.1.

We tested PCA group separation using PERMANOVA (*vegan::adonis2)* on Euclidean distances. For independent group comparisons, we used Wilcoxon rank-sum tests; for one-sample comparisons, we used Wilcoxon signed-rank or Fisher’s exact test as appropriate. For motif and gene-set enrichment analyses, we used a Benjamini-Hochberg FDR correction. We performed all analyses in R unless otherwise specified.

### Data Availability

### Code Availability

Add Github ID for code availability.

## Results

### Promoter methylation distinguishes between sporadic IPF and control subjects

To identify changes in DNA methylation between control (declined, age-matched donor; male n=5, female n=5) and IPF (explant lungs; male n=7, female n=7) subjects (**Supplemental Table 1**), we performed long-read DNA sequencing on CD326+ lung epithelium (**Fig. 1A**). Subjects were matched for age (median control 65 (IQR 61-68), median IPF 66.5 (IQR 63-70)), with 3 control and 7 IPF subjects reporting smoking history (**Supplemental Table 1B**). The mean genome coverage per subject ranged from 27 to 58 (median 30; **Supplemental Table 2**). Approximately 27.3 million CpGs, of the approximately 28 million represented in the hg19 reference genome^30^, met our ≥ 5x coverage threshold in ≥ 9 control and ≥ 12 IPF subjects and were included in downstream analysis.

**Figure 1:**
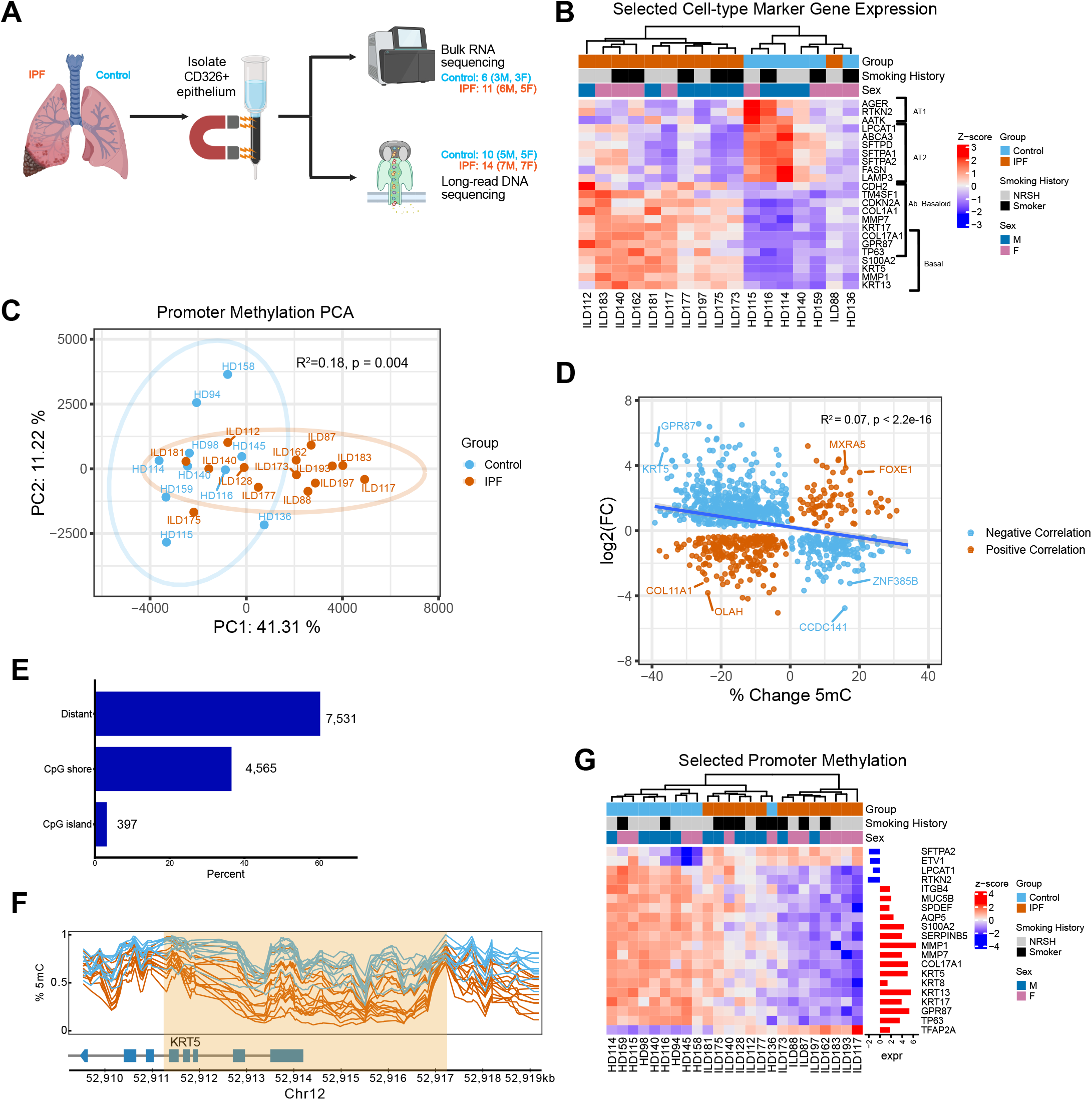
Promoter methylation distinguishes between IPF and control patients. **A** Schematic representing the study design. We isolated CD326+ epithelium from IPF and control donors and performed long-read sequencing (Oxford Nanopore Technologies) and bulk RNA-sequencing to identify DNA methylation changes and associated transcriptional alterations. **B** Unsupervised hierarchical clustering of subjects based on row-scaled expression of AT1, AT2, aberrant basaloid, and basal marker genes. The genes associated with each cell type are shown in brackets. **C** PCA based on DNA methylation at promoter tiles. PCA was performed after mean imputation of missing values by region. Patients with IPF are shown in vermillion and control subjects are shown in blue. Significance assessed by PERMANOVA on the PCA distance matrix. **D** Correlation between the percent change in methylation at differentially methylated promoters and log_2_(fold change expression) of the associated gene. Positively correlated associations are shown in vermillion and negatively correlated associations are shown in blue. Significance assessed by simple linear regression. **E** The distribution of the proximity of differentially methylated promoter tiles to CpG islands. The number of sites for each category is shown next to the respective bar. **F** *KRT5* genomic track showing a differentially methylated promoter. Vermillion lines indicate IPF subjects and blue lines indicate control subjects. The percentage of methylated cytosines out of total cytosines sequenced is plotted by genomic coordinate. The yellow highlighted region represents twelve consecutive tiles that were significantly differentially methylated between groups. **G** Unsupervised hierarchical clustering of subjects based on row-scaled methylation of differentially methylated promoters. The log_2_(fold change expression) in IPF relative to control epithelium is shown for each associated gene.

To investigate the relationship between methylation and transcription, we performed bulk RNA-sequencing on a subset of subjects with available samples (control male n=3, female n=3; IPF male n=6, female n=5; **Fig. 1A**). 25,075 genes had detectable expression in ≥ 5 control or ≥ 9 IPF subjects and were included in further analysis. Analysis of our RNA-sequencing data revealed a total of 1,308 differentially expressed genes (DEGs; log_2_(fold change expression) ≥ 1, q-value ≤ 0.05; **Extended Data Table 1**). IPF and control groups exhibited modest clustering in PCA space based on gene expression (R^2^=0.19, p=0.006; **Supplemental Fig. S1A**). The contribution of sex (median 2.2%) and reported smoking history to individual feature variance was negligible (median 2.8%; **Extended Data Table 2**).

We then sought to identify the differences in transcriptional signatures of the epithelial cell populations isolated from IPF and donor lungs. AT1 markers, including *AGER, RTKN2,* and *AATK*, and AT2 markers, including *LPCAT1, ABCA3, SFTPD, SFTPA1, SFTPA2, FASN,* and *LAMP3*, were significantly downregulated in IPF relative to control EPCAM+ cells (**Fig. 1B**). KRT5-/KRT17+ cells markers, including *CDH2, TM4SF1, CDKN2A, COL1A1, MMP7,* and *KRT17* and basal markers, including COL17A1, GPR87, TP63, *S100A2, KRT5, MMP1,* and *KRT13*, were significantly upregulated in IPF relative to control EPCAM+ cells (**Fig. 1B**). These data are consistent with a loss of the alveolar epithelium and an increase in basal and other proximal airway cell-types in the IPF distal lung tissue samples, similar to other reports^4–6^.

As promoter (defined here as 3000 bp surrounding the TSS) methylation is canonically associated with repression of gene activity, we first characterized methylation at these loci^31,32^. The methylation percentages of individual CpGs at each promoter region were aggregated across 500 bp windows and used for downstream analysis. IPF and control groups exhibited modest separation in PCA space based on DNA methylation of promoter tiles found on autosomal chromosomes, with disease state explaining 18% of variance by PERMANOVA, p=0.004 (**Fig. 1C**). Sex accounted for a median of 4.8% of the overall variance across individual features, with 97.2% of regions with a contribution to variance greater than 50% found on the X-chromosome (**Extended Data Table 3**). The contribution of reported smoking history to variance was negligible (1.8%; **Extended Data Table 3**).

Differential methylation analysis revealed a total of 12,493 DMRs (6,831 unique genes; defined as FDR-adjusted p-value ≤ 0.01 and change of ≥ 10% in methylation; **Extended Data Table 4**). To investigate the relationship between promoter methylation and associated gene expression, we intersected DMRs identified in all subjects with differentially expressed genes identified in subjects with RNA-sequencing data (**Fig. 1D**). We observed a modest, but significant, negative correlation between changes in DMR methylation and DEG expression between control and IPF subjects (Simple-linear regression, R^2^=0.07, p<2.2 x 10^-16^), consistent with the canonical relationship between promoter methylation and gene expression^9^. Of the 1,010 total associations, 690 were negatively correlated, and 320 were positively correlated. Overall, genes with promoter DMRs were significantly enriched for differential expression (Fisher’s exact test, p < 2.2^-16^, OR = 2.09).

The majority of DMRs were found inside CpG-island (CGI) shores (within 2000 bp of CGI; **Fig. 1E**). This finding is consistent with the results of methylation mapping studies that have shown that hypomethylated promoters (particularly those overlapping CpG-dinucleotide enriched CpG islands) typically have stable patterns of methylation with differentiation and across cell types^31–34^.

Among the most significant promoter DMRs with corresponding changes in expression, we identified the AT2 marker genes *SFTPA2* and *ETV1* and genes associated with aberrant epithelial differentiation (e.g. *ITGB4*, *MUC5B, S100A2, KRT5, and KRT17*) (**Fig. 1F&G**)^4–6^.

Interestingly, while promoter methylation of the AT2 marker gene *LPCAT1* and the AT1 marker gene *RTKN2* was decreased, these genes were downregulated in IPF relative to control subjects. Similarly, the promoter DMR at the TFAP2A locus was hypermethylated, with gene expression increased, in IPF subjects. These noncanonical relationships between promoter methylation and gene activity may reflect silencer activity of these loci or regulation by a TF that preferentially binds methylated DNA^35–37^.

These results are consistent with an expansion of KRT5-/KRT17+ cells and a loss of alveolar epithelial in the IPF distal lung. The modest correlation between respective gene promoter DNA methylation and gene expression suggests interaction between promoter DNA methylation and additional layers of gene regulation.

### DNA methylation variability is enriched at putative distal regulatory elements

As changes in methylation at regions distal to TSSs have been shown to account for the majority of variation across somatic cell types and disease states^12,13,15,38,39^, we broadened our analysis to encompass the entire genome. To identify putative regulatory elements and assess consistent patterns of DNA methylation across consecutive CpGs, we used *MethyLasso* to perform region-based segmentation and methylation state classification (**Fig. 2A**). These regions ranged in length from 30 to 47,482 bp (median 262 bp) (**Supplemental Fig. S2A**), with no difference in region size observed between disease states (Wilcoxon rank-sum test p-value = 0.92).

**Figure 2:**
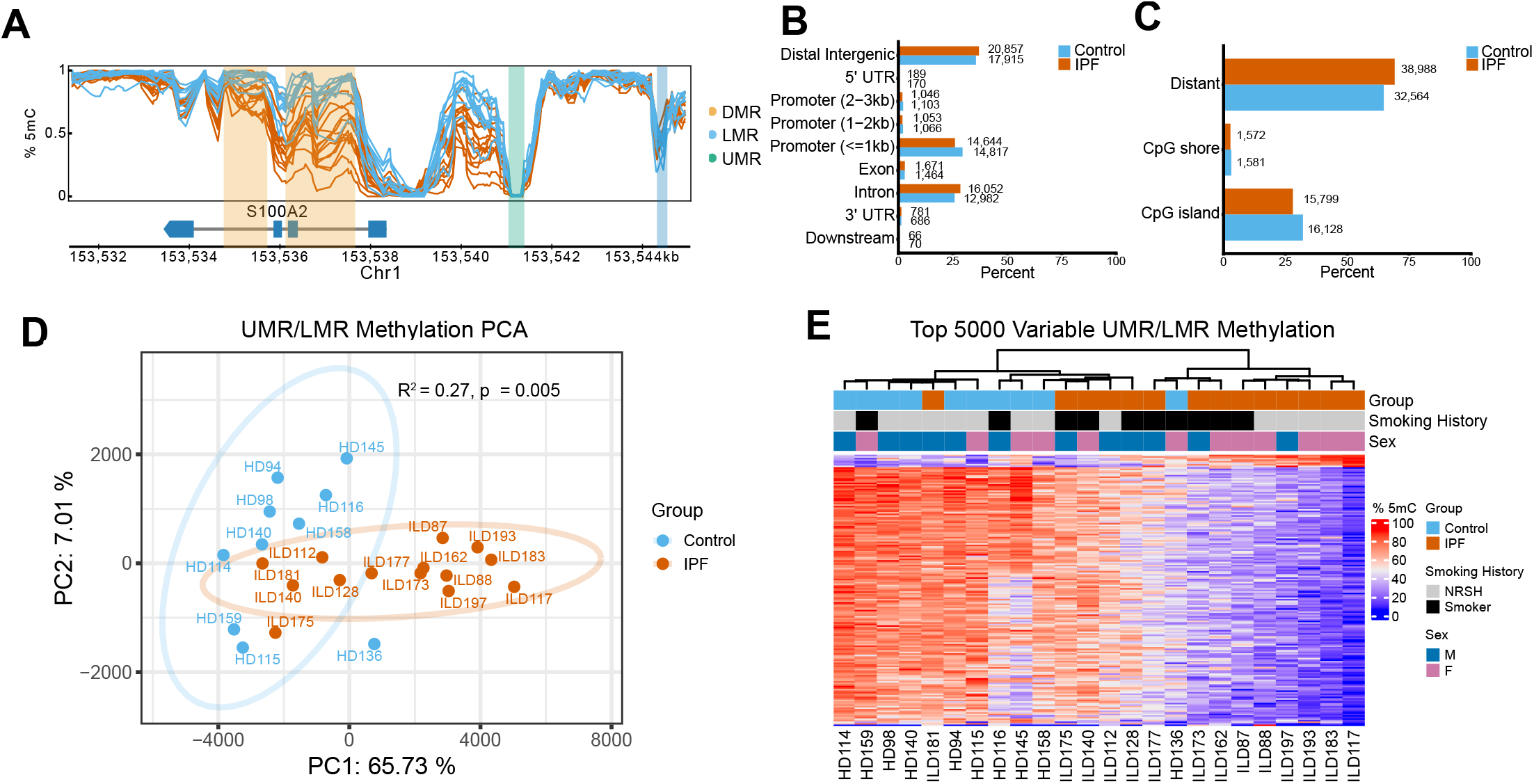
DNA methylation variability is enriched at putative distal regulatory elements. **A** S100A2 genomic track showing the results of genome-wide segmentation. A UMR is shown in green, an LMR is shown in blue, and a DMR is shown in yellow. **B** Distribution of the genomic region annotations of UMRs and LMRs by group. The number of sites for each category is shown next to the respective bar. **C** Distribution of the proximity UMRs and LMRs to CpG islands by group. **D** PCA based on the methylation percentages at UMRs and LMRs on autosomal chromosomes. PCA was performed after mean imputation of missing values by region. Patients with IPF are shown in vermillion and control subjects are shown in blue. Significance assessed by PERMANOVA on the PCA distance matrix. **E** Unsupervised hierarchical clustering of subjects based on DNA methylation of the top 5,000 most variable UMRs and LMRs (excluding those present on the X-chromosome) across all samples.

The emergence of UMRs and LMRs is associated with cell differentiation and has been shown to denote regulatory elements^12,13,15,38^. To perform unbiased analysis of the methylation landscapes of IPF and control epithelium at putative regulatory regions, we annotated UMRs and LMRs in both groups separately. We then merged annotations from both groups, identifying a total of 27,667 UMRs and 20,816 LMRs common to both groups. 548 UMRs and 6,983 LMRs were unique to control subjects, and 1,989 UMRs and 18,143 LMRs were unique to IPF subjects (**Extended Data Table 5**).

We next sought to describe the proximity of UMRs and LMRs identified across groups to annotated genomic elements. Using the R package *TxDB.Hsapiens.UCSC.hg19.knownGene*, we assigned UMRs and LMRs to the nearest-neighbor TSS (**Extended Data Table 5**). Approximately 35% of UMRs and LMRs were localized to a distal intergenic region, 30% in promoters, and 25% in intronic regions (**Fig. 2B**). UMRs and LMRs were largely distal to CGIs (**Fig. 2C**). Consistent with previous studies, the majority (76%) of stably hypomethylated promoter regions were found at CGIs^18,34^ (**Extended Data Table 5**).

To assess disease-associated variability at putative regulatory elements identified across the genome, we evaluated methylation at all UMRs and LMRs identified across control and IPF subjects. IPF and control groups clustered separately in PCA space based on UMR and LMR methylation, with disease state explaining 27% of methylation variance by PERMANOVA, (p=0.003), a larger variance than was seen in promoter focused analysis (**Fig. 2D**). Relative to region frequency, introns, exons, 3’ and 5’ UTRs and distal intergenic regions exhibited a greater contribution to the first principal component than promoter sites (**Supplemental Fig. S2B**). Sex accounted for a median of 7% of the overall variance, with 96% of regions with a contribution to variance greater than 50% found on the X-chromosome. The contribution of reported smoking history to variance was negligible (2%; **Extended Data Table 6**). Unsupervised hierarchical clustering of the 5,000 most variable regions on autosomal chromosomes revealed distinct patterns of UMR and LMR methylation in control and IPF groups (**Fig. 2E**).

These findings show that UMR/LMRs identified across the genome display genomic distributions and methylation states consistent with putative regulatory elements and account for a larger proportion of differentially methylated regions associated with IPF than analysis focused solely on promoter regions.

### Genome-wide DMRs are predominantly hypomethylated in the IPF epithelium and enriched for association with TFs implicated in aberrant epithelial differentiation

As the methylation states of putative regulatory elements identified in the control and IPF epithelium were variable across disease states, we sought to characterize regions of significant differential methylation and their proximity to potential drivers of aberrant epithelial differentiation. Using *MethyLasso*, we identified 61,760 DMRs (defined as FDR-adjusted p-value ≤ 0.01 and absolute change in methylation ≥ 10%; 14,442 unique genes; **Extended Data Table 7**) out of 1,202,124 tested regions. Consistent with our promoter-restricted analysis, the majority of DMRs reflected hypomethylation in IPF relative to control (52,059 hypomethylated and 9,701 hypermethylated; **Fig. 3A** and **Supplemental Fig. 3A**). DMRs ranged in length from 8 to 6,413 bp (median 349 bp; **Supplemental Fig. S2C**).

**Figure 3:**
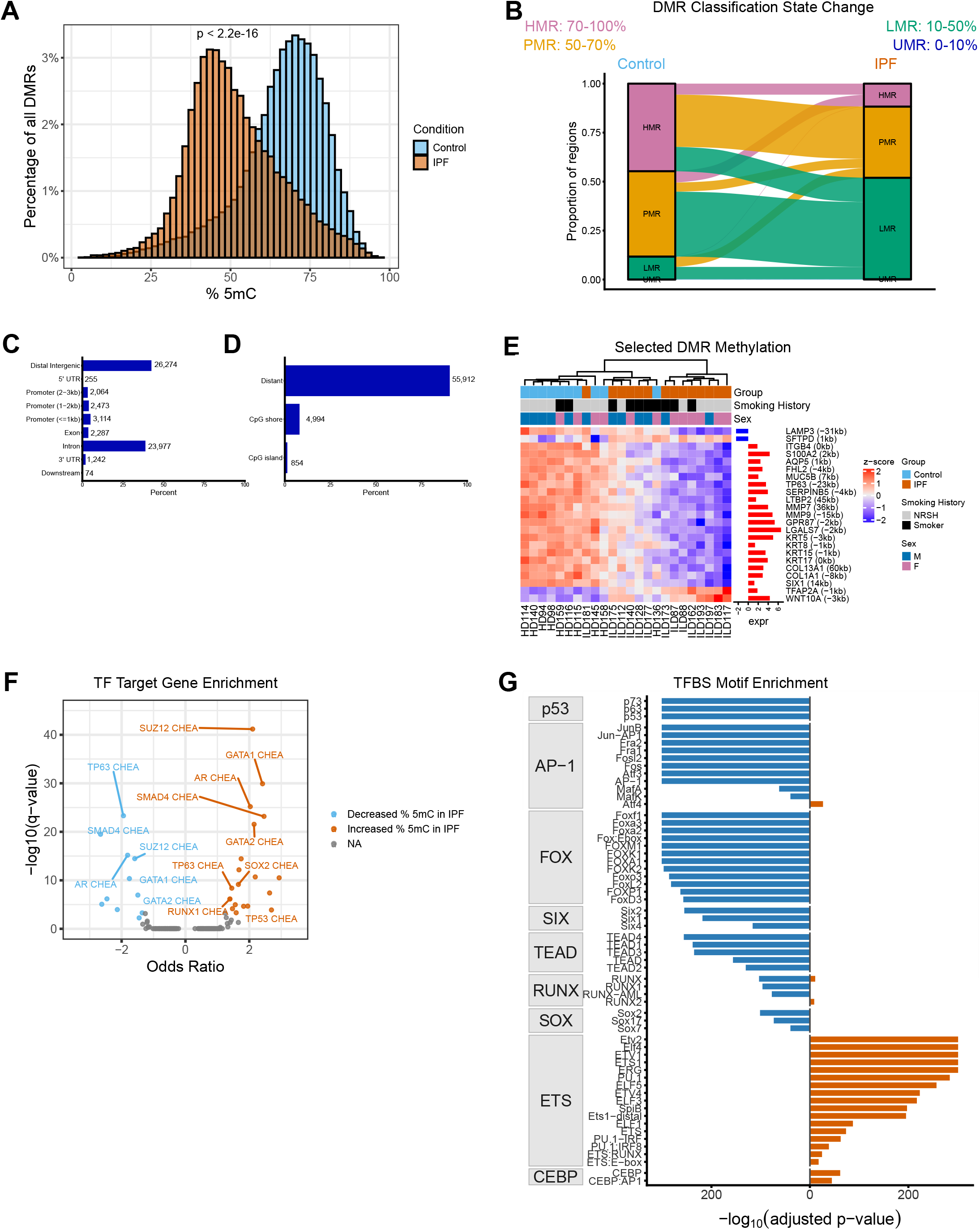
DMRs are enriched for association with TFs implicated in aberrant epithelial differentiation. **A** Distribution of methylation percentages at DMRs by group. The median methylation was 49% for IPF and 68% for control subjects. Significance assessed by wilcoxon rank-sum test. **B** Change in methylation classification (HMR, PMR, LMR, UMR) at DMRs between control and IPF groups. HMRs are shown in pink, PMRs are shown in yellow, LMRs are shown in green, and UMRs are shown in blue. **C** Distribution of the proximity of DMRs to genomic region annotations. The number of sites for each category is shown next to the respective bar. **D** Distribution of the proximity of DMRs to CpG islands. **E** Unsupervised hierarchical clustering of subjects based on methylation of DMRs. The log_2_(fold change expression) in IPF relative to control epithelium is shown for each associated gene. **F** Gene set enrichment analysis based on genes associated with hypo- or hypermethylated DMRs in IPF. The -log_10_(q-value) is plotted against the odds ratio for each enriched TF. The enriched TFs were identified from a total of 104 terms included in the *ENCODE_and_ChEA_Consensus_TFs_from_ChIP-X* library. **G** Transcription factor binding sequence motif enrichment analysis based on genomic coordinates of DMRs with hypo- or hypermethylation in IPF. The -log_10_(q-value) is shown for each enriched TFBS motif. TFs are grouped by family. The enriched TFs were identified from a total of 472 motifs included in the *homer2* known motifs library.

The difference in the distribution of methylation values in IPF relative to control at DMRs was highly significant (Wilcoxon rank-sum test, p-value < 2.2^-16^). On average, IPF subjects had 51% methylation at DMRs, whereas control subjects had 66% methylation (median of 49% and 68% for IPF and control, respectively). To understand patterns of change in DNA methylation at DMRs in IPF relative to control subjects, we classified methylation states in both groups. The predominant change in classification was a shift from a PMR to an LMR (20,454 sites; **Fig. 3B**). In contrast, the change in classification observed in the fewest sites was that of a shift from a UMR to an LMR (14 sites). A similar predominance of DNA hypomethylation was observed at promoters (**Supplemental Fig. 3B**).

The majority of DMRs were intronic or intergenic (approximately 40% each), with 12% localized to promoter regions (**Fig. 3C**). These regions were largely distal to CGIs (**Fig. 3D**). Annotation with the nearest TSS using *TxDB.Hsapiens.UCSC.hg19.knownGene* revealed an association of DMRs with 14,422 unique genes.

To understand how shifts in methylation relate to altered pathway activity in the IPF epithelium, we investigated significant DMRs and their nearest-neighbor gene associations (**Fig. 3E**). DMRs were identified at regions proximal to AT2-marker genes downregulated in IPF, including *SFTPD*, a surfactant protein^40^, and *LAMP3*, a lysosomal membrane protein involved in surfactant homeostasis^41^. Among upregulated genes, we identified significant alterations in methylation at sites proximal to *MUC5B*, a gene with expression typically restricted to the airway epithelium that has increased activity in the distal lung in IPF^42^ and basal cell markers *TP63*, *GPR87, KRT5* and *KRT17*.

To identify transcriptional programs potentially associated with the differential methylation observed in the IPF epithelium, we performed gene set enrichment analysis using *enrichR*. DMRs were grouped by hypo- or hypermethylation, and the single nearest neighbor gene was assigned and used as input for gene set enrichment analysis. TF target gene enrichment was largely similar across hypo- and hypermethylated DMR associations, consistent with the context-dependent relationship between DNA methylation and gene activity. We identified significant enrichment of p63, a master regulator of basal cell identity that has been implicated in aberrant distal epithelial remodeling in IPF^4,6^ and FOSL2, an AP-1 complex subunit involved in airway secretory cell fate transition to AT2s following injury^43^(**Fig. 3F** and **Supplemental Fig. S3C**). Further, target genes for SUZ12, a component of the PRC2 complex shown to be necessary for maintaining AT2 fate^44,45^, were significantly enriched in IPF DMRs (**Fig. 3F** and **Supplemental Fig. S3C**). SMAD4 target gene enrichment, a mediator of pro-fibrotic TGFβ signaling, were associated with both hypo- and hypermethylated DMRs^46,47^ Consistent with the results of our DMR TF association enrichment analysis, SUZ12, SMAD4, and p63 target gene expression was upregulated in the IPF epithelium (**Supplemental Fig. S3D**).

To further assess TF binding site enrichment within DMRs, exploring the relevance of these regions as regulatory elements, we performed transcription factor binding site (TFBS) motif enrichment analysis using *homer2* (**Fig. 3G** and **Supplemental Fig. S3E**). While this approach allows identification of motifs characteristic of TF families, the high motif similarity across members of the same family does not allow prioritization of individual TFs within a family. TFBS motif enrichment analysis of both promoter-restricted and genome-wide regions revealed enrichment of p53 (family includes p63 and p73), forkhead box (FOX), AP-1, TEAD, and SIX family motifs among hypomethylated promoter DMRs (**Fig. 3, Supplemental Fig. 3D**). FOX family proteins are key regulators of alveolar epithelial function; FOXA2 is required for epithelial maturation,^48^ whereas FOXM1 has been implicated in pro-fibrotic activation and proliferation of the lung epithelium^49^. TEAD family TFs act downstream of Hippo-YAP/TAZ signaling, a pathway involved in alveolar epithelial differentiation and repair processes that is aberrantly upregulated in the IPF epithelium^50–58^. *SIX1* is a developmental TF involved in early lung morphogenesis and is reactivated in the IPF epithelium^59^. In contrast, among hypermethylated regions, we observed enrichment of ETS-domain TFs (**Supplemental Fig. S3E**). ETS domain-containing transcription factors, including ETS1 and ELK1, have been implicated in pro-fibrotic activation of the lung epithelium^60,61^.

In addition to motifs enriched in promoter DMRs, in DMRs identified using genome-wide segmentation, we also observed enrichment of RUNX and SOX family TFs in hypomethylated regions and CEBP family TFs in hypermethylated regions (**Fig. 3G**). RUNX2 has been implicated in the profibrotic activation of AT2 cells in the IPF epithelium and has been shown to interact with YAP/TAZ^55,62–64^. SOX2 is required for airway epithelial cell proliferation and SOX2+ airway epithelium migrates to the distal lung in response to injury^65,66^. CEBPA is required for AT2 fate maintenance and loss of the activity of this TF drives lung fibrosis^67,68^.

Here, we demonstrate a predominance of hypomethylation at genome-wide DMRs in the IPF epithelium that associates with both transcriptional changes of genes and predicted differential activity of TFs implicated in failed adaptive epithelial repair processes.

### DMR methylation is associated with expression of disease-associated genes

To identify expression quantitative trait methylation (eQTMs), we used the R package *Matrix eQTL* to find significant correlations between methylation of DMRs and expression levels of DEGs (**Extended Data Table 8**). For this analysis, only potential cis-regulatory relationships were targeted, and therefore, a maximum threshold of 1 million base-pairs between the TSS of the DEG and the midpoint of the DMR was used^9^. Of the 69,604 total comparisons, 39,196 were significant (q-value ≤ 0.1; **Fig. 4A**). The identified associations were modestly, but significantly, biased toward the negative, consistent with the canonical negative association between methylation and expression (median *β* = -0.023; one-sided exact binomial test, p < 2.2^-16^).

**Figure 4:**
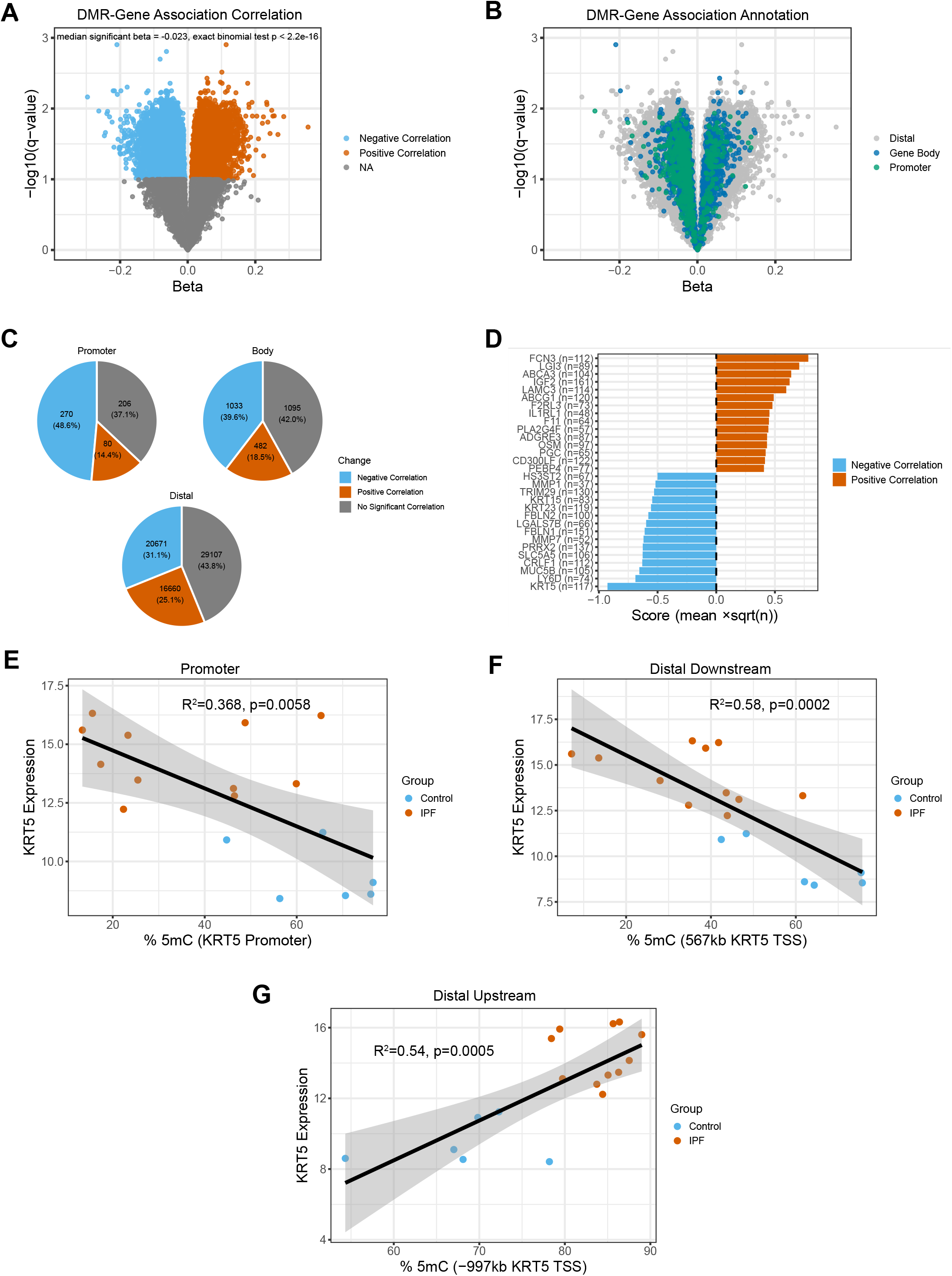
DMR methylation shows significant correlation with transcriptional activity of DEGs. **A** eQTM analysis results showing significant (FDR-adjusted p-value ≤ 0.1) negative correlations in blue, and positive correlations in vermillion. The -log_10_(q - value) is plotted against the beta value for each DMR-gene expression comparison. Significance of negative correlation bias assessed by exact binomial test. **B** eQTM analysis results showing promoter DMRs (within 3000 bp of a TSS, green), gene-body associated DMRs (between +3000 bp of a TSS and the end of the associated gene, blue), and distal DMRs (outside of the promoter and gene body of the associated gene, grey). **C** Proportion of eQTMs by proximity to the associated gene. **D** Top 15 positively and negatively correlated eQTMs ranked by number of associated DMRs for each gene and average effect size of correlation. **E** Correlation between variance-stabilized, normalized *KRT5* expression and methylation at a promoter DMR. Significance assessed by simple linear regression. **F** Correlation between variance-stabilized, normalized *KRT5* expression and methylation at a DMR 567 kb downstream of the *KRT5* TSS. Significance assessed by simple linear regression. **G** Correlation between variance-stabilized, normalized *KRT5* expression and methylation at a DMR -997 kb upstream of the *KRT5* TSS. Significance assessed by simple linear regression.

To understand how the location of the DMR might affect the relationship between methylation and expression, we annotated DMRs according to presence in the associated gene’s body, promoter, or outside of either region (distal; **Fig. 4B**). In DMRs found within promoters, most associations were consistent with the expected negative correlation between methylation and expression (**Fig. 4C**). However, for DMRs found in gene bodies or at regions distal to the associated gene, the predominance of a negatively correlated relationship was less marked (**Fig. 4C**).

We further characterized these relationships by ranking significant associations according to the product of their mean beta value and the square root of the number of associated DMRs (**Fig. 4D**). Among the top 15 positively and negatively correlated genes were basal-cell marker genes *KRT5* and *KRT15* (**Fig. 4E-G**), and *MUC5B,* an airway-secretory marker with ectopic expression in IPF. Interestingly, the expression of AT2 marker gene *ABCA3* (a phospholipid transporter) was positively correlated with DMR methylation changes. Expression of *ABCA3* is decreased in the IPF epithelium, reflecting a loss of AT2 cells in disease. The relationships identified by the eQTM analysis indicate significant correlations between DMR hypomethylation and *ABCA3* downregulation in IPF.

We performed a second eQTM analysis targeting relationships between promoter DMRs and DEGs (**Extended Data Table 8**). This revealed a total of 966 significant associations out of 1,501 total tests (**Supplemental Fig. S4A**).The majority of these associations reflected a negative correlation between methylation and expression (median *β* = -0.038; one-sided exact binomial test, p < 2.2^-16^; **Supplemental Fig. S4B**). Among the top 15 negatively correlated genes, the majority of promoter tiles for basal-cell marker genes *KRT5, KRT15,* and *KRT17* were differentially methylated and significantly associated with transcriptional activity (**Supplemental Fig. S4C**). *ZNF385B* had the most significant correlation between promoter methylation and DEG expression (**Supplemental Fig. S4D**). This gene has not been extensively characterized in IPF, but is significantly downregulated in the lung epithelium in our RNA-sequencing data and is suggested to be a biomarker for breast and ovarian cancer ^69,70^. In summary, our eQTM analysis reveals significant correlations between numerous intergenic, intragenic, and promoter DMRs and DEGs implicated in aberrant epithelial differentiation.

### Loss of alveolar epithelial fate is associated with epigenetic repression of lineage specific transcription factor binding sites

As the heterogeneity of a CD326+ epithelial cell selection strategy and the bulk nature of our assay does not allow us to determine cell-type specificity for these DNA methylation changes, we sought to validate these results by assessing changes in DNA methylation during in vitro trans-differentiation of AT2 cells, the primary epithelial stem cell of the lung alveolus. To drive AT2-to-basal cell metaplastic trans-differentiation, 3D co-culture of HTII-280+ AT2 with primary human mesenchyme was used as previously described, a model in which AT2 cells transdifferentiate into basal and airway-like epithelium after 14-days of culture and recapitulates some of the aberrant features of the IPF distal lung^8^. We isolated HTII-280 cells from four age-matched, non-diseased control lungs (N = 2 male and N=2 female; **Supplemental Table 3**) and performed differential methylation analysis between freshly isolated HTII-280+ AT2 cells (Day 0) and all CD326+ epithelial cells isolated from organoids after 14-days of co-culture with human mesenchymal cells(Day 14) (**Fig. 5A**).

**Figure 5:**
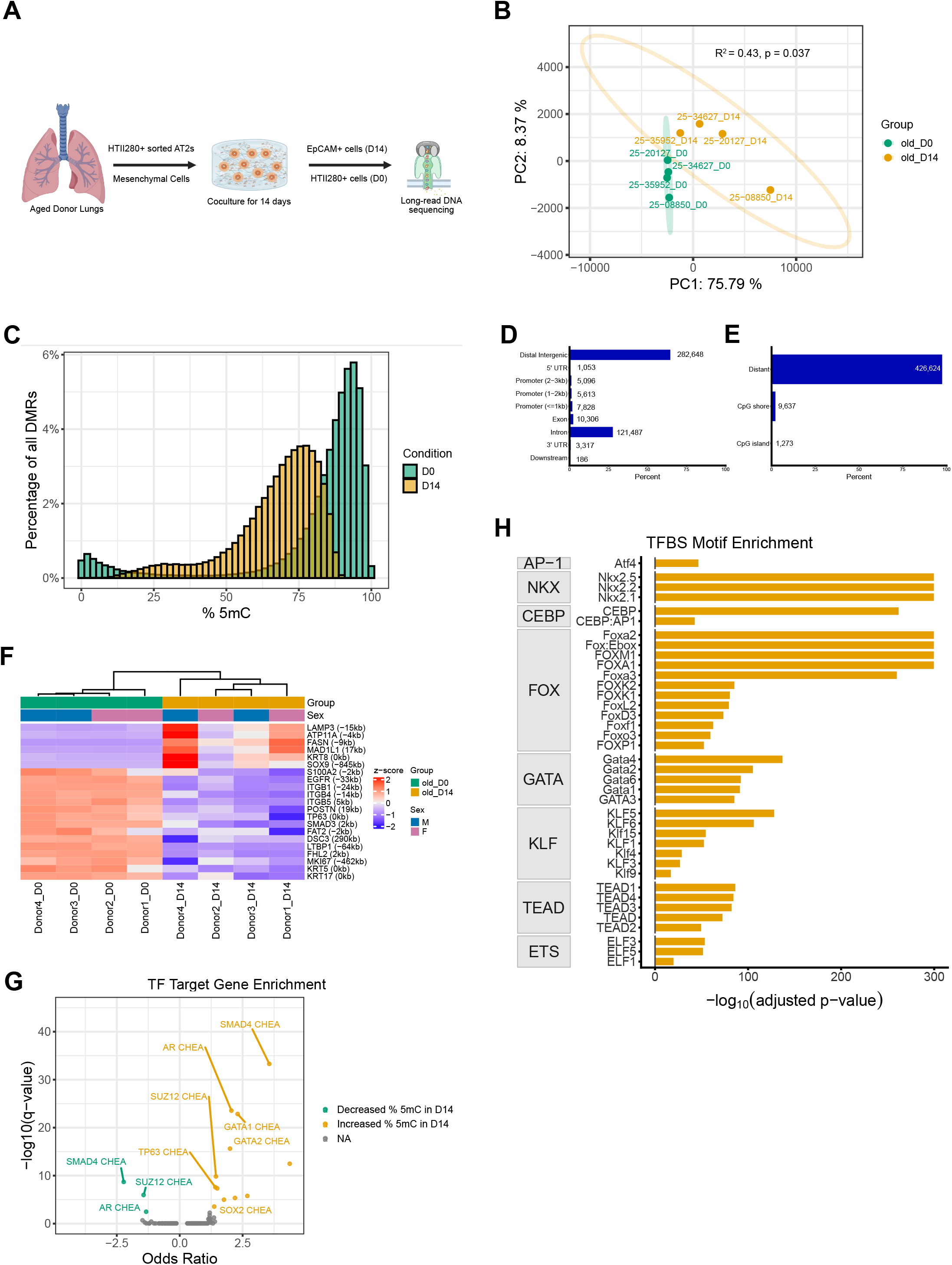
Loss of alveolar epithelial fate is associated with epigenetic repression of lineage specific transcription factor binding sites. **A** Schematic representing the experimental design. We isolated HTII280+ AT2s from aged donors and performed long-read sequencing (Oxford Nanopore Technologies) on samples pre- and post-co-culture with mesenchymal cells. **B** PCA based on the methylation percentages at UMRs and LMRs on autosomal chromosomes. PCA was performed after mean imputation of missing values by region. Day 0 samples are shown in green and D14 samples are shown in yellow. Significance assessed by PERMANOVA on the PCA distance matrix. **C** Distribution of methylation percentages at DMRs by condition. The median methylation was 71% for D14 and 89% for D0 samples. Significance assessed by Wilcoxon rank-sum test. **D** Distribution of the proximity of DMRs to genomic region annotations. The number of sites for each category is shown next to the respective bar. **E** Distribution of the proximity of DMRs to CpG islands. **F** Unsupervised hierarchical clustering of subjects based on row-scaled methylation of the most significant DMR associated with the labeled gene. The distance of the DMR from the TSS is shown for each gene. **G** Gene set enrichment analysis based on genes associated with hypo- or hypermethylated DMRs in D14 samples. The -log_10_(q-value) is plotted against the odds ratio for each enriched TF. The enriched TFs were identified from a total of 104 terms included in the *ENCODE_and_ChEA_Consensus_TFs_from_ChIP-X* library. **H** Transcription factor binding sequence motif enrichment analysis based on genomic coordinates of DMRs with hypo- or hypermethylation associated with AT2 transdifferentiation. The -log_10_(q-value) is shown for each enriched TFBS motif. The TF family is shown for each enriched TF. The enriched TFs were identified from a total of 472 motifs included in the *homer2* known motifs library.

Day 0 and Day 14 samples clustered separately in PCA space based on UMR and LMR methylation (R^2^=0.44, p=0.037; **Fig. 5B**). Consistent with patterns of differential methylation observed between IPF and control epithelium, post-co-culture samples reflected a predominance of hypomethylation at DMRs relative to Day 0 pre-co-culture samples (p < 2.2^-16^; **Fig. 5C**). We identified a remarkable number of differentially methylated regions in this model, 437,534 total DMRs between day 0 and day 14, with 39,882 becoming hypermethylated and 397,652 becoming hypomethylated during in vitro culture (18,327 unique genes; **Extended Data Table 9**). These DMRs were largely found at distal intergenic and intronic regions (65% and 28%) and distal to CGIs (98%; **Fig. 5D&E**).

We identified significant hypermethylation at multiple regions proximal to AT2 marker genes *LAMP3*, *ATP11A,* and *FASN* and hypomethylation at regions proximal to basal cell marker and pro-fibrotic activation associated genes, including *KRT5, KRT17, TP63, LTBP1, ITGB4,* and *S100A2* (**Fig. 5F**). Interestingly, the promoter of the alveolar transitional cell marker *KRT8* was hypermethylated in post-co-culture samples, potentially reflecting the presence of a silencer element at this locus.

TF target gene enrichment analysis revealed enrichment of target genes for TFs including SMAD4, SUZ12, and AR at DMRs (**Fig. 5G**). In contrast with previous enrichment analyses, p63 target genes were significantly enriched for association with hypermethylated regions only.

TFBS motif analysis revealed significant enrichment of NKX2, CEBP, and FOX family motifs among hypermethylated regions (**Fig. 5H**). NKX2.1 is a lung epithelium lineage TF involved in maintenance of alveolar epithelial identity and regulation of AT2 to AT1 differentiation^53,71,72^. Interestingly, no TFBS motifs were significantly enriched among our hypomethylated set. This is potentially due to the large number of hypomethylated regions tested, or the fact that we are comparing homogeneous HTII-280+ AT2 cells to a heterogeneous cell population after 14-days of culture.

In summary, the AT2 aberrant differentiation that occurs following co-culture with mesenchymal cells is associated with widespread changes in DNA methylation, including hypermethylation at predicted binding sites of TFs necessary for maintaining AT2 identity, including CEBP and NKX2, which are likely driving the phenotypic loss of alveolar cell fate seen in vitro.

### Shared IPF- and AT2 co-culture-associated hypomethylated regions are enriched for p53 and AP-1 family motifs

To identify changes in methylation that were reproducible across IPF epithelium and experimental AT2 transdifferentiation, we filtered for (“consensus”) DMRs with 50% reciprocal overlap in both systems (**Extended Data Table 11**). Across the total 10,101 overlapping DMRs, the percentages of change in methylation were poorly correlated (simple linear regression R^2^ = 0.01, p < 2.2^-16^; **Fig. 6A**), likely reflecting cell type and system dependent methylation changes. Despite this heterogeneity, the majority of overlapping regions, 5,983, reflected hypomethylation in both systems and exhibited a strong correlation (Pearson r = 0.44, FDR = 3.3^−288^; **Fig. 6B**).

**Figure 6:**
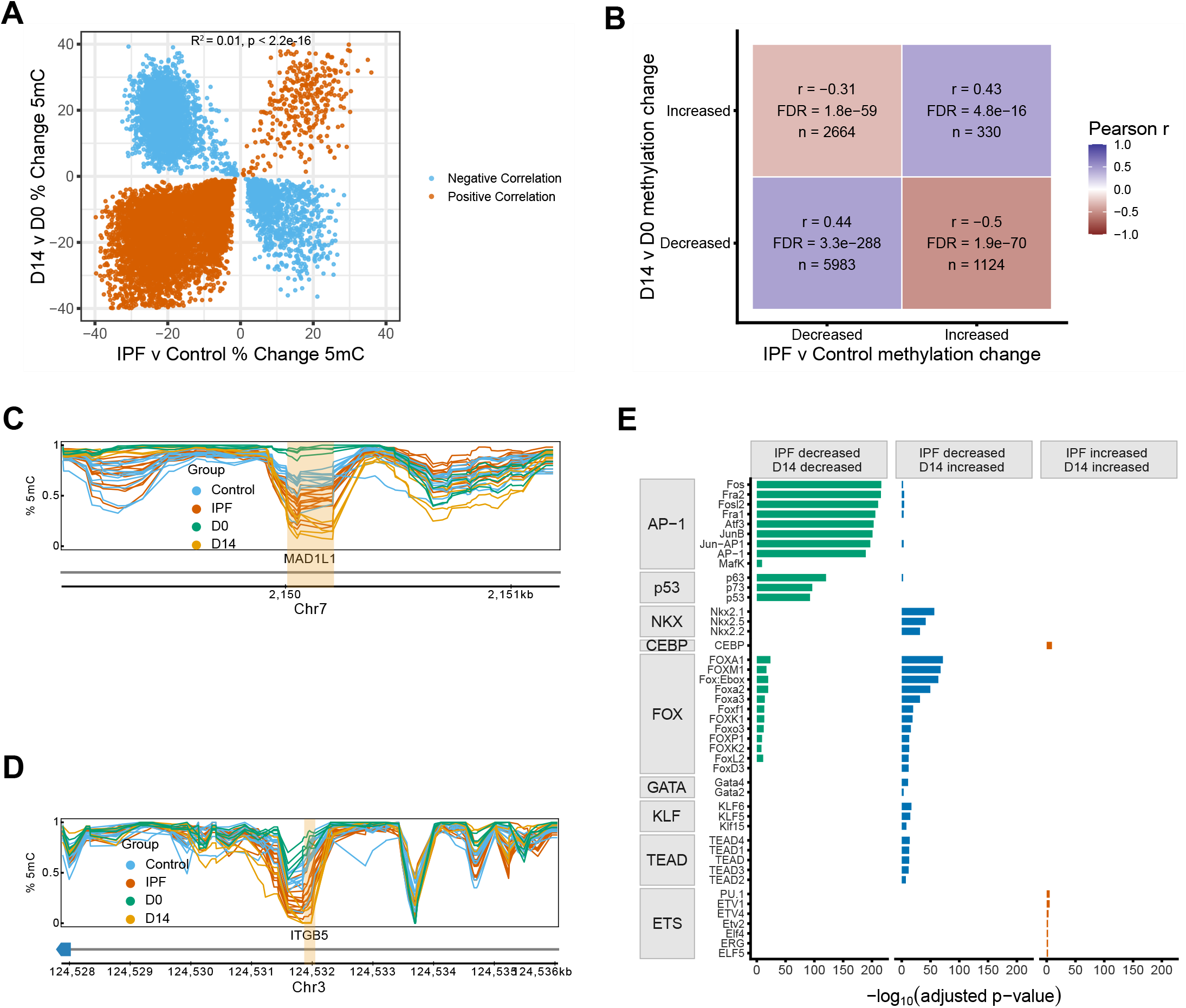
Shared IPF- and AT2 co-culture-associated hypomethylated regions are enriched for p53 and AP-1 family motifs. **A** Correlation between percentage change in methylation for one-to-one overlapping DMRs identified IPF versus control subjects and D14 versus D0 AT2 co-culture conditions. Significance was assessed by simple linear regression. **B** Pearson correlation between IPF-associated and D14-associated percentage change in methylation after stratifying overlapping DMRs by direction of methylation change. Tile labels show Pearson r and the number of overlapping DMR pairs by class. **C** *MAD1L1* genomic track showing region hypomethylated in IPF and following mesenchyme co-culture in yellow. **D** *ITGB5* genomic track showing region hypomethylated in IPF and following mesenchyme co-culture in yellow. **E** Transcription factor binding site motif enrichment analysis of one-to-one overlapping DMRs stratified by methylation-change direction. The -log_10_(q-value) is shown for each enriched TFBS motif. The TF family is shown for each enriched TF. The enriched TFs were identified from a total of 472 motifs included in the *homer2* known motifs library.

Focusing on this subset, we identified consensus hypomethylated regions that included sites intronic to *ITGB5* and *MAD1L1* (**Fig. 6C&D**). *MAD1L1* encodes a component of the mitotic-spindle assembly checkpoint and is a susceptibility locus for IPF. *ITGB5,* as well as another consensus hypomethylated region-associated gene *ITGB8* (**Extended Data 10**), are upregulated in the airway basal cells of IPF patients^73^, and function in TGF-*β* activation and signaling^74,75^.

We next sought to identify if consensus DMRs were associated with common transcriptional regulatory programs. Although there was no enrichment across all hypomethylated regions associated with experimental AT2 transdifferentiation, consensus hypomethylated regions were significantly enriched for multiple p53 and AP-1 family members (**Fig. 6E**). Consensus hypermethylated regions, on the other hand, showed only modest enrichment of ETS and CEBP family TFs (**Fig. 6E**). Interestingly, regions hypermethylated with AT2 transdifferentiation and hypomethylated in the IPF epithelium were significantly enriched for NKX2 and FOX family motifs (**Fig. 6E**), potentially reflecting loss of alveolar epithelial lineage fidelity in the co-culture model and convergence of diverse epithelial populations toward an indeterminate epithelial cell state in the IPF lung.

Overall, overlapping DMRs show a predominance of hypomethylation with enrichment of TFBS motifs for AP-1 and p53 TF families, suggesting that these transcription factors play a key role during both in vitro and in vivo transdifferentiation of alveolar lineages.

### IPF GWAS variants overlap DMRs

IPF GWAS have revealed that the majority of disease-associated variants reside outside of coding regions^43^. We hypothesized that overlap between single-nucleotide polymorphisms (SNPs) and DMRs in the IPF epithelium or AT2s following mesenchyme co-culture would identify candidate loci with regulatory roles in aberrant alveolar repair and fibrotic progression, predicting roles for these SNP-DMR overlaps. Filtering for direct overlap of SNPs identified by a recent IPF GWAS^76^ and IPF epithelial DMRs revealed a total of 90 unique, disease-associated variants (Bonferroni-adjusted significance threshold of 1.96^-9^; **Fig. 7A**; **Extended Data Table 11**). An additional 356 SNPs were found overlapping DMRs associated with AT2 transdifferentiation (**Supplemental Fig. S6A**; **Extended Data Table 11**). Similar to the full set of disease-associated SNPs, SNPs overlapping DMRs identified in either system were largely intergenic and intronic (**Fig. 7B & Supplemental Fig. S6B**).

**Figure 7:**
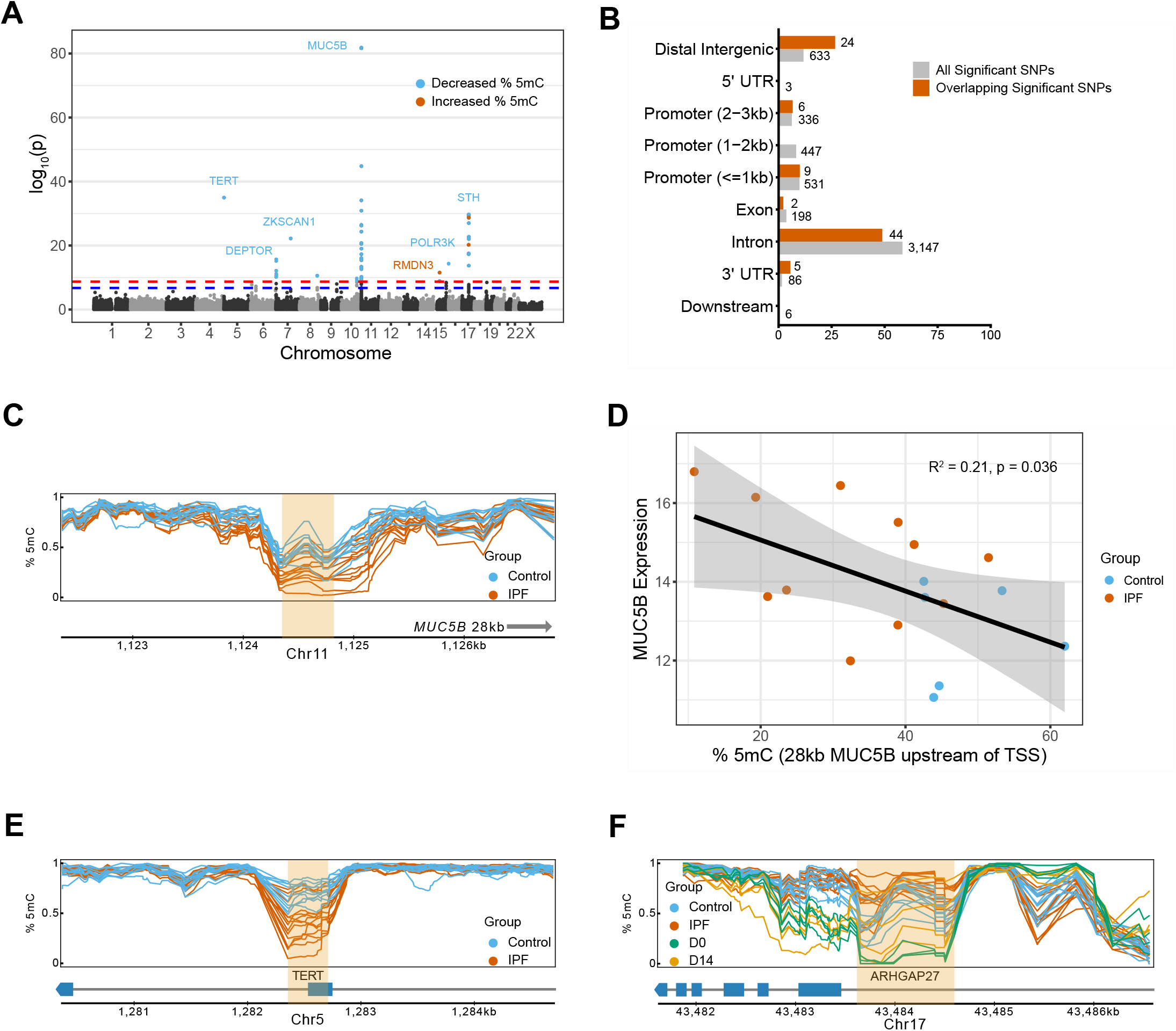
IPF GWAS variants overlap DMRs. **A** Location of SNPs with direct overlap with a DMR. SNPs associated with DMRs with decreases in methylation are shown in blue, and those associated with DMRs with increased methylation are shown in vermillion. **B** Distribution of the proximity of SNPs to genomic region annotations. The number of sites for each category is shown next to the respective bar. Annotations for all significant SNPs are shown in gray and annotations for significant SNPs overlapping a DMR are shown in vermillion. **C** Intergenic genomic track with distance to nearest-neighbor gene *MUC5B* labeled. DMR overlapping a significant disease-associated variant is shown in yellow. **D** Correlation between variance-stabilized, normalized *MUC5B* expression and methylation at a DMR 36kb upstream of the *MUC5B* TSS. **E** *TERT* genomic track showing DMR overlapping a significant disease-associated variant in yellow. **E** *ARHGAP27* genomic track showing region hypermethylated in IPF and following mesenchyme co-culture overlapping a significant disease-associated variant in yellow.

DMRs overlapped some of the most significant GWAS loci, including regions proximal to the *MUC5B* locus (**Fig. 7C & Supplemental Fig. S6C**). *MUC5B* has been extensively studied in IPF pathogenesis, with evidence suggesting that ectopic expression of *MUC5B* in the alveolar epithelium may predispose individuals to failed repair through increased endoplasmic reticulum stress^77,78^. Methylation of the IPF epithelial DMR overlapping the *MUC5B-*proximal SNP rs57614263 was significantly correlated with *MUC5B* expression (simple linear regression R^2^ = 0.415, p = 0.0017; **Fig. 7D**), consistent with a regulatory role for this element. We also identified the *MUC5B* proximal variant rs12802931 within a DMR associated with AT2 transdifferentiation (**Supplemental Fig. S6C**). This variant is in linkage disequilibrium with the *MUC5B* promoter polymorphism, rs35705950^79^, and therefore does not represent an independent signal. However, the association of hypomethylation at this locus with aberrant alveolar epithelial differentiation suggests a regulatory role for this site. We further identified differential methylation at the rs7725218 locus (**Fig. 7E**). This variant is intronic to *TERT,* a gene that encodes the catalytic component of telomerase, deficits in which are predicted to promote early telomere shortening^80^.

Finally, direct overlap between DMRs identified in IPF versus control subjects and D14 versus D0 AT2 co-culture conditions, revealed hypermethylation of an intronic region of *ARHGAP27* containing two disease associated variants, rs62064603 and rs73984391 (**Fig. 7F**). Although neither variant has been studied in the context of IPF, the concordance between differential methylation at this site across both systems supports a regulatory role for this locus.

Our GWAS-DMR overlap analysis reveals colocalization of numerous SNPs with DMRs associated with the IPF epithelium or AT2 transdifferentiation, including loci proximal to *MUC5B* and *TERT*, highlighting putative regulatory elements that may contribute to aberrant epithelial differentiation.

## Discussion

While previous studies of DNA methylation in IPF have largely relied on array-based analyses of heterogeneous whole lung tissue^9,81–83^, our study provides high-resolution, genome-wide characterization of CpG methylation in the lung epithelium. To our knowledge, this study represents the first whole-genome DNA methylation analysis of IPF subjects. Using a sliding-window promoter-restricted analysis, we identified 12,493 differentially methylated tiles associated with 6,831 unique genes. We showed that, using a genome-wide segmentation approach, we were able to more accurately discriminate between control and IPF groups and identify intergenic and intronic disease-associated methylation alterations. This method allowed for the identification of 61,760 DMRs associated with 14,422 unique genes. Overall, DMRs were generally localized to intergenic and intronic regions and found distal to CGIs. Top DMRs were identified at regions proximal to basal-cell marker genes *KRT5* and *GPR87* and the secretory cell marker *MUC5B*. TF enrichment analysis consistently predicted increased activity of p63 at hypomethylated DMRs. eQTM analysis showed 35,555 significant associations between DMR methylation and DEG expression, including *FASN, ABCA3, KRT5, KRT15, MMP7,* and *MUC5B*, genes implicated in failed repair and fibrotic activation^4–6^. DMRs overlapped GWAS loci, including *TERT* and *MUC5B-*associated regions. Methylation analysis of an experimental model of disease-associated AT2 transdifferentiation revealed widespread changes in DNA methylation and enrichment of CEBP and NKX2 family motifs among hypermethylated regions. DMRs identified in both systems were largely hypomethylated and enriched for p53 (including p63/p73) and AP-1 family motifs.

In a developmental context, hypomethylated regions distal to TSSs emerge with cell-type specification and often correspond to cis-regulatory elements enriched for lineage-determining TF binding and gene associations^12–15,84^. Our hypothesis that DMRs present in the IPF epithelium denote regulatory elements responsible for the alterations in transcriptional activity that drive failed repair and the progression of fibrotic disease is supported by the highly significant enrichment of TF motifs in the p53 family including p63, which has been implicated in aberrant epithelial activity and repair processes, among DMRs identified in the IPF epithelium and consensus IPF epithelium/AT2 transdifferentiation regions. Reactivation of p63 has been reported in KRT5+ basal cells that transdifferentiate from AT2s co-cultured with IPF mesenchyme^8^ and p63 expression has also been shown to localize to intermediate alveolar epithelial cells found in fibrotic regions of the lung^85^. p63 target genes are also among the most highly enriched in upregulated genes identified by our RNA-sequencing data.

Several TF families implicated in epithelial injury responses and aberrant repair are enriched for association with hypomethylated regions identified in this study. Members of the AP-1 complex, including FOSL2, showed significant gene association and binding site motif enrichment among hypomethylated regions in the IPF epithelium. Following severe injury, distal airway resident secretory (SCGB1A1+) cells have been shown to mobilize to the alveoli and differentiate into alveolar epithelium^86,87^. FOSL2/FRA2 activity, downstream of NOTCH inhibition, has been shown to be necessary for and to mediate this process^43^. Additionally, a recent study of chromatin accessibility in the IPF epithelium revealed highly significant enrichment of AP-1 family TF motifs among regions with increased accessibility in KRT5-/KRT17+ cells^88^.The described differential methylation in this study at these loci may reflect a historical record of AP-1 activity in the IPF epithelium^12,13,38,89–92^.

A second pathway instrumental in AT2 and AT1 differentiation during development and in response to injury is HIPPO-YAP/TAZ, which regulates transcriptional activity through the TEAD family of TFs^50,51,93^. While YAP/TAZ signaling is necessary for AT2 to AT1 differentiation, increased activity of this pathway has been observed in the IPF epithelium, and mechanistic studies have shown that sustained activity drives aberrant differentiation^55–57^. Consistent with our data, previous studies in human cancer cell lines have shown that most YAP1 and TEAD binding sites are distal to TSSs at putative enhancer elements associated with transcriptional regulation of YAP-target genes^93^. Physical interactions between YAP/TAZ and RUNX2 have been described to modulate expression of genes involved in growth and differentiation^63^. Previous work from our group has shown that sustained activation of YAP/TAZ results in increased chromatin accessibility at the RUNX2 promoter in AT2s from bleomycin injured mice^55^. Other groups have shown increased expression of RUNX2 in profibrotic AT2s, with knockdown of this TF impairing their proliferation^64^.

A second class of methylation changes associated with pathways implicated in early developmental processes and lineage fidelity was observed in this study. Chromatin analysis of alveolar epithelial differentiation has suggested activity of FOXA1, in conjunction with NKX2.1, may regulate epigenomic remodeling during the AT2 to AT1 transition^71^. Indeed, physical interaction of NKX2.1 and FOXA1 has been reported in the lung epithelium, although the relationship was one of inhibition^94^. NKX2.1 has also been shown to be necessary, through binding to separate regions, for maintenance of both AT2 and AT1 identity^72^. Furthermore, loss of NKX2.1 drives loss of AT2 identity and aberrant differentiation in alveolar epithelial progenitor population^95^. Consistent with this, our data show significant enrichment of NKX2 family motifs among regions hypermethylated in an experimental model of AT2 transdifferentiation. These data support a role for epigenetic repression through DNA methylation in driving loss of AT2 identity and the emergence of an aberrant, basal-like phenotype.

The FOX family TFBS motif enrichment signal was present across DMRs present in IPF epithelium and AT2 transdifferentiation analyses, albeit in opposite directions. FOXM1 is necessary for profibrotic activation of the alveolar epithelium in response to TGF*β* stimulation^49^. In a murine model, FOXM1 activation in AT2s exacerbates fibrotic remodeling by activating the SNAIL1 promoter and increasing the inflammatory response^96^. Additionally, AT2-specific FOXM1 deletion impairs proliferation and differentiation^97^. FOXA1/2 have been shown to be necessary for alveolarization and secretory cell development^48,98–100^. FOXA1 motifs specifically are enriched at regions of open chromatin that emerge with AT2 to AT1 differentiation^71^. Additionally, FOX family members including FOXK2 and FOXP1 have been shown to increase WNT/β-catenin signaling^101^. WNT signaling governs respiratory airway secretory cells (RASC; SCGB3A2+) to AT2 cell differentiation^102^. An expansion of RASC-derived AT2s in the distal lung is consistent with the patterns of DNA methylation observed in our data. As epigenomic profiling of AT2 to AT1 differentiation has shown colocalization of FOXA1 with NKX2.1 motifs, our data may reflect diverging, context-dependent roles for FOXA1, or other members of this TF family, in aberrant, disease-associated differentiation^71^. Alternatively, the observed enrichment of FOX motifs at regions hypermethylated in our AT2 transdifferentiation model may emerge as a consequence of culture conditions.

Consistent with a role for DNA methylation in the epigenetic repression of lineage maintaining TFBS in IPF, we identify hypermethylation of CEBPA binding sites in our data. CEBP family member CEBPA is an AT2 specific TF involved in maintenance of AT2 identity^67^. Further, a mutant form of CEBPA has been shown to directly interact with and inhibit the de novo methyltransferase DNMT3A, suggesting a potential role for CEBPA and epigenetic regulation^103^. Our group’s prior work demonstrated that aberrant sustained activation of YAP/TAZ-TEAD activity in mouse AT2 cells resulted in loss of CEBPA expression leading to aberrant epithelial cell differentiation following bleomycin injury^55^.

In addition to predicted alterations at loci associated with specific transcriptional pathways, we observed further methylome dysregulation at regions associated with PRC2, a chromatin remodeling complex involved in widespread transcriptional repression that serves to establish and maintain the repressive H3K27me histone mark^104,105^. The enrichment of EZH2 and a second member of PRC2, SUZ12, among both hyper- and hypo-methylated DMRs in IPF epithelium and our experimental AT2 transdifferentiation model, and upregulated SUZ12 target DEGs in the IPF epithelium, is suggestive of dysregulated PRC2 activity in aberrant epithelial differentiation. While a recent study has shown that PRC2 activity is necessary to maintain AT2 fate^44^, other research has implicated noncanonical activity of EZH2 in aberrant epithelial differentiation^45^. Given the presence of PRC2 component target gene enrichment across the modalities and systems investigated in this study, this complex represents a promising target for further investigation of epigenetic regulation of failed epithelial repair in IPF.

Together, our motif and target gene enrichment analyses identify regulatory programs associated with DMRs in the IPF epithelium; the eQTM analysis further links these changes to dysregulation of specific genes implicated in failed repair processes. While the results of our eQTM analysis indicate a bias toward negatively correlated DMR-DEG relationships, a substantial number of the identified associations reflect positively correlated relationships. A potential regulatory relationship for all of these associations is inconsistent with the canonical repressive role for methylation at regulatory elements. While the associated DMR loci are potentially silencer elements or CTCF insulators at topologically associating domains (TAD) boundaries, an alternative hypothesis is that these associations do not reflect regulatory relationships with genes such as *ABCA3* and *FASN*, but instead the emergence of hypomethylation at regulatory elements that are associated with disease-emergent cell types that appear concomitant with the loss of the alveolar epithelium. The expression of these marker genes is lost in IPF and is significantly correlated with the decrease in methylation at the associated DMRs. Overall, the strong correlation (both positive and negative) between disease-associated transcriptional dysregulation and DMR demethylation is suggestive of a role for the identified DMRs in IPF progression.

In addition to their relationship with transcriptional dysregulation, these DMRs may provide context for the functional role for numerous IPF-susceptibility loci. Here, we show that a total of 90 IPF-associated SNPs overlap DMRs in the IPF epithelium, while 356 overlap DMRs associated with AT2 transdifferentiation. Among these variants, the most significant mapped to loci upstream of *MUC5B* and intronic to *TERT.* In CD326+ epithelium, the expression of *TERT* does not differ appreciably between groups, and the impact of hypomethylation at this locus in IPF subjects is unclear and warrants further investigation. In contrast, the *MUC5B* proximal SNP is found within a DMR with a significant loss of methylation in IPF and a significantly correlated increase in MUC5B expression. Although many of the sites identified by our GWAS-DMR overlap analysis are not the lead variants at their respective loci, their localization to regions with significant alterations in methylation in either system is highly suggestive of a potential regulatory role. These data suggest that variants that may have been deprioritized because of linkage disequilibrium with stronger association signals, may still highlight functional regulatory regions.

An important limitation of our study is the expected heterogeneity of cell-type composition of a CD326+ epithelial selection from distal lung tissue. Thus, although we identify a transcriptional signature of the loss of alveolar epithelium along with the increase in basal and KRT5-/KRT17+ cells populations in our IPF subjects, we are unable to definitively attribute the observed loss of DNA methylation in the IPF epithelium to aberrant alveolar epithelial differentiation. Alternatively, the loss of methylation in IPF may reflect the persistence of hypomethylation at regulatory elements specific to the airway epithelium as these cells migrate to the distal lung, acquiring a methylation signature of the alveolar epithelium. Further investigation of epithelial subtype-specific methylation signatures could be used to identify whether the observed trends in methylation reflect migration of distal airway cells or the aberrant differentiation of the alveolar epithelium. While the selection method we employed in our study complicates interpretation, a recent study suggests that the widely-used alternative method of AT2 isolation, HTII-280 selection, does not allow for purification of AT2 cells from the IPF lung^106^. Instead, an HTII-280 sort from fibrotic human lungs results in a heterogeneous population of basal, club, goblet, and terminal and respiratory bronchiolar secretory cells. As DNA methylation serves as a historical record of cell lineage, and a comparison of a relatively homogenous AT2 population from control lungs with a heterogeneous alveolar and airway epithelial population from the IPF lung could bias regions of differential methylation toward those representing airway lineage rather than features of aberrant differentiation, we believe the CD326+ selection to remain the optimal choice for our study.

In summary, our study represents the first whole-genome DNA methylation analysis conducted in the lung epithelium of IPF subjects. We show a predominance of DNA hypomethylation in the IPF epithelium localized to regions associated with p63 activity, as well as other TFs implicated in pro-fibrotic activation and maintenance of alveolar epithelial fate. We further demonstrate that AT2 transdifferentiation is associated with a similar predominance of hypomethylation; however, hypermethylation occurs at predicted binding sites for TFs necessary for maintaining lineage fidelity, which are repressed as a result. Overall, the majority of DMRs were distal to promoter regions, showing a concentration in intergenic and intronic regions. Based on these results, we propose a model wherein changes in DNA methylation at distal regulatory elements is concomitant with and facilitates increases in transcriptional activity of genes involved in failed epithelial repair, aberrant differentiation, and loss of lineage fidelity in the lung epithelium. The putative distal regulatory elements, particularly those overlapping GWAS loci at regions proximal to *MUC5B, TERT,* and *ARHGAP27*, identified by this study as well as the TFs predicted to act in regions of altered DNA methylation, merit further functional and mechanistic study.

**Supplemental Figure 1:**
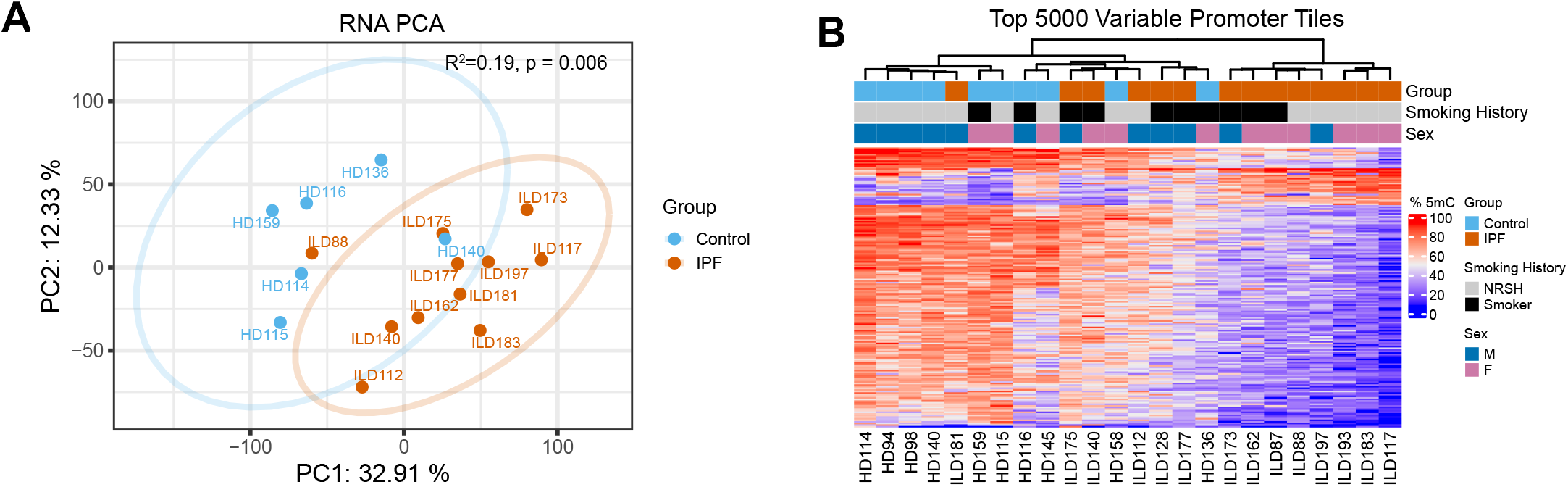
Promoter methylation distinguishes between IPF and control patients. **A** PCA based on variance-stabilized, normalized gene expression. Patients with IPF are shown in vermillion and control subjects are shown in blue. Significance assessed by PERMANOVA on the PCA distance matrix. **B** Unsupervised hierarchical clustering of subjects based on methylation percentages of the top 5,000 most variable promoter tiles (excluding those present on the X-chromosome).

**Supplemental Figure 2:**
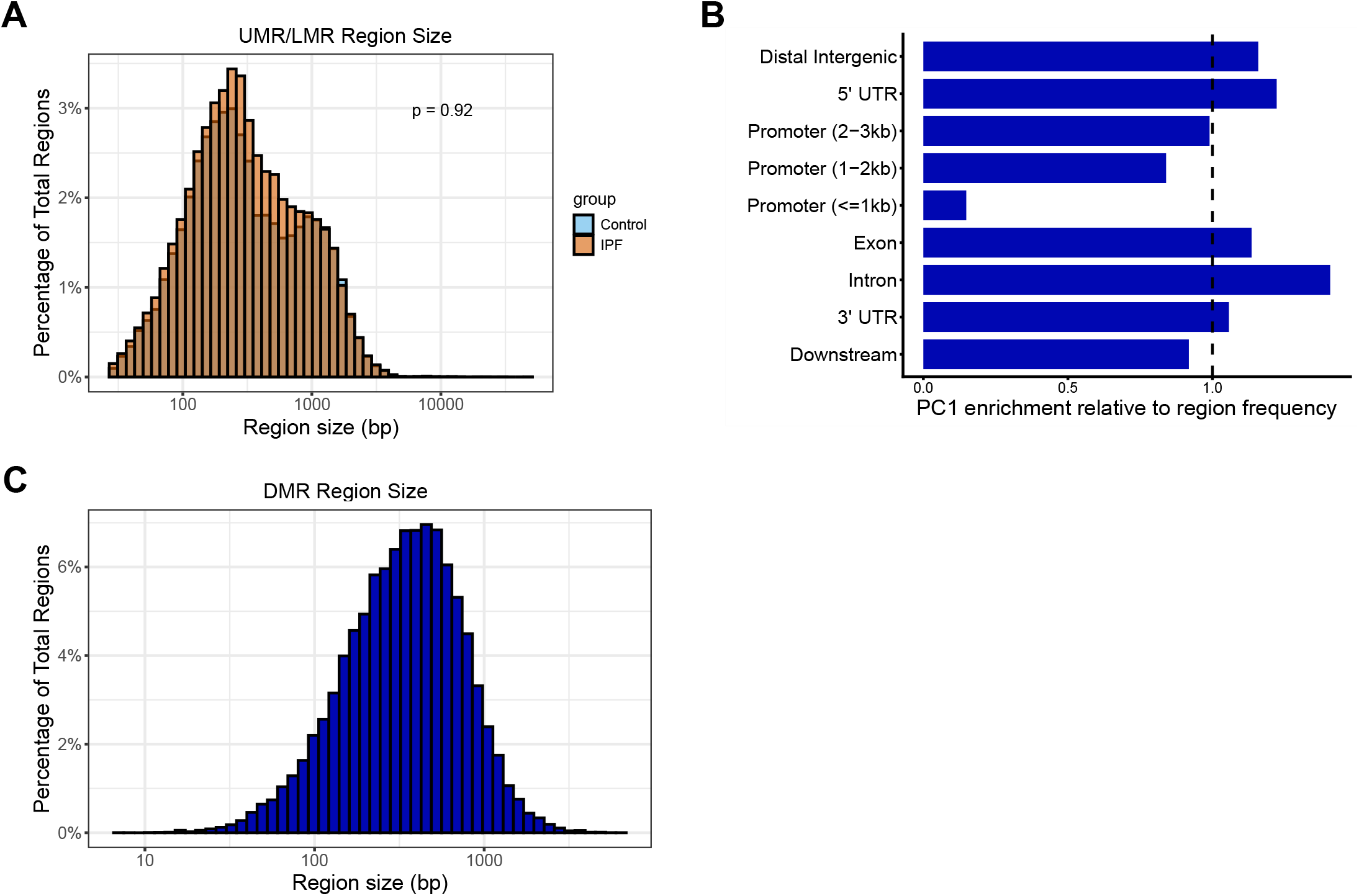
DNA methylation variability is enriched at putative distal regulatory elements. **A** Distribution of UMR and LMR region-size in base pairs by group. Significance assessed by wilcoxon rank-sum test. **B** Contribution of genomic annotations to PC1. The enrichment was calculated as the fraction of summed squared PC loadings within an annotation class divided by the fraction of regions assigned to the class. **C** Distribution of region-size in base pairs of DMRs.

**Supplemental Figure 3:**
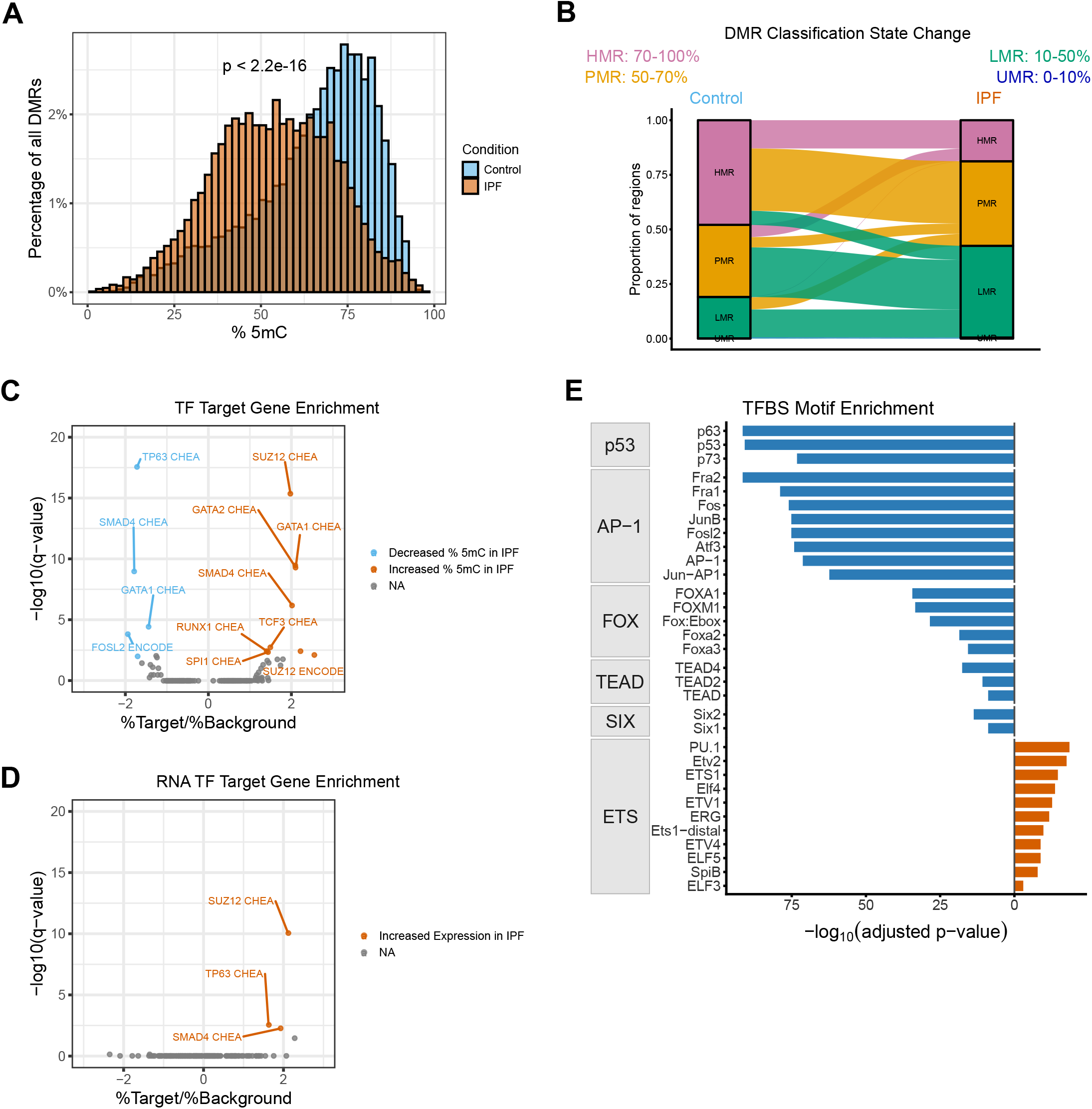
Hypomethylated promoters are enriched for association with p63 in IPF. **A** Distribution of methylation percentage at differentially methylated promoter tiles by group. IPF methylation percentages are shown in vermillion, and control methylation percentages are shown in blue. The median methylation was 54% for IPF and 69% for control subjects. Significance assessed by wilcoxon rank-sum test. **B** Change in methylation classification (HMR, PMR, LMR, UMR) at differentially methylated promoters between control and IPF groups. HMRs are shown in pink, PMRs are shown in yellow, LMRs are shown in green, and UMRs are shown in blue. **C** Gene set enrichment analysis based on genes associated with hypo- or hypermethylated promoters in IPF. The -log_10_(q-value) is plotted against the odds ratio for each enriched TF. The enriched TFs were identified from a total of 104 terms included in the *ENCODE_and_ChEA_Consensus_TFs_from_ChIP-X* library. **D** Gene set enrichment analysis based on upregulated and downregulated genes in IPF. The -log_10_(q-value) is plotted against the odds ratio for each enriched TF. The enriched TFs were identified from a total of 104 terms included in the *ENCODE_and_ChEA_Consensus_TFs_from_ChIP-X* library. **E** Transcription factor binding site motif enrichment analysis based on genomic coordinates of promoter tiles with hypo- or hypermethylation in IPF. The -log_10_(q-value) is shown for each enriched TFBS motif. TFs are grouped by family. The enriched TFs were identified from a total of 472 motifs included in the *homer2* known motifs library.

**Supplemental Figure 4:**
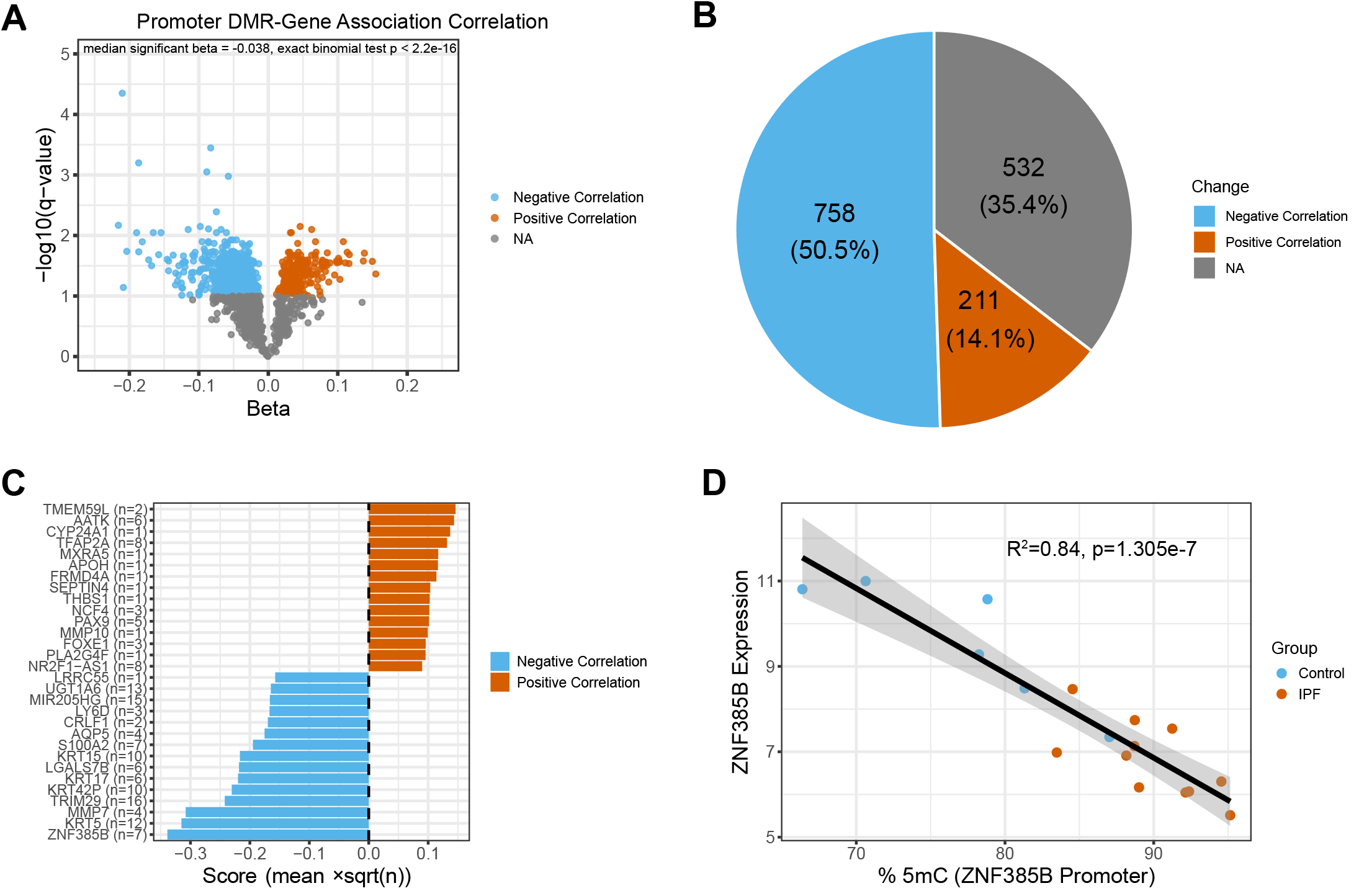
Promoter DMR methylation shows a significant correlation with transcriptional activity of DEGs. **A** eQTM analysis results of differentially methylated promoters identified by tiling, showing significant (FDR-adjusted p-value ≤ 0.1) negative correlations in blue, and positive correlations in vermillion. The -log_10_(q - value) is plotted against the beta value for each DMR-gene expression comparison. Significance of negative correlation bias assessed by exact binomial test. **B** Proportion of DMR-gene expression comparisons with significant correlations with gene expression. **C** Top 15 positively and negatively correlated eQTMs ranked by number of associated DMRs for each gene and average effect size of correlation. **D** Correlation between variance-stabilized, normalized *ZNF385B* expression and methylation at a promoter DMR. Significance assessed by simple linear regression.

**Supplemental Figure 5:**
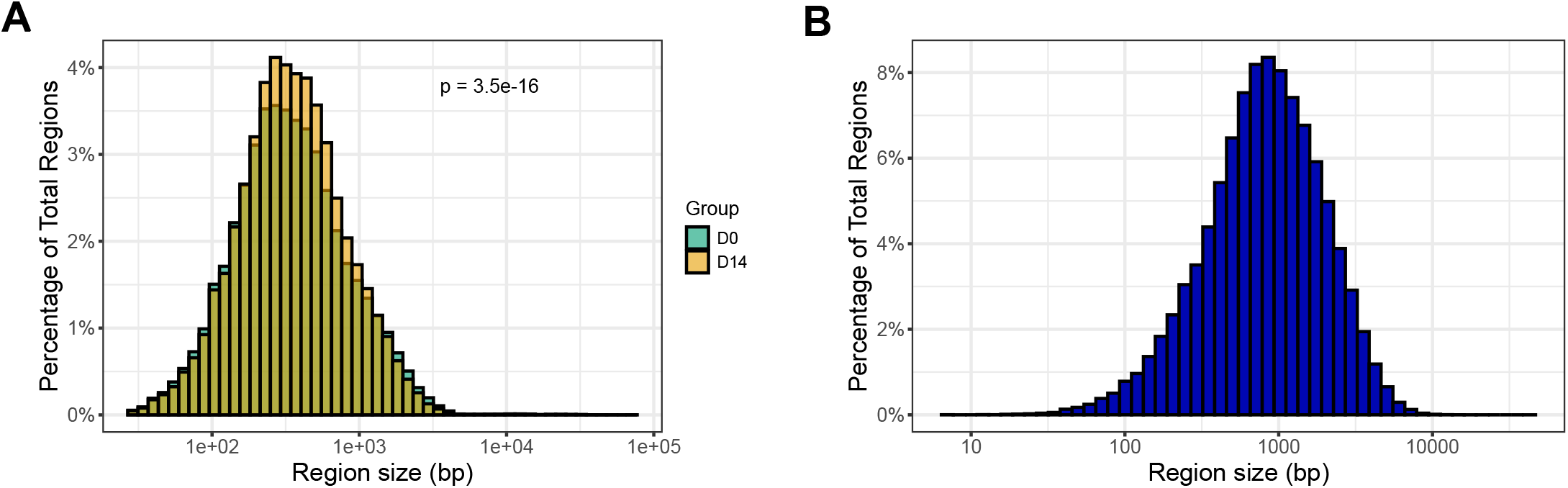
Disease-associated variants are enriched in DMRs. **A** Distribution of UMR and LMR region-size in base pairs by group. Significance assessed by wilcoxon rank-sum test. **B** Distribution of region-size in base pairs of DMRs.

**Supplemental Figure 6:**
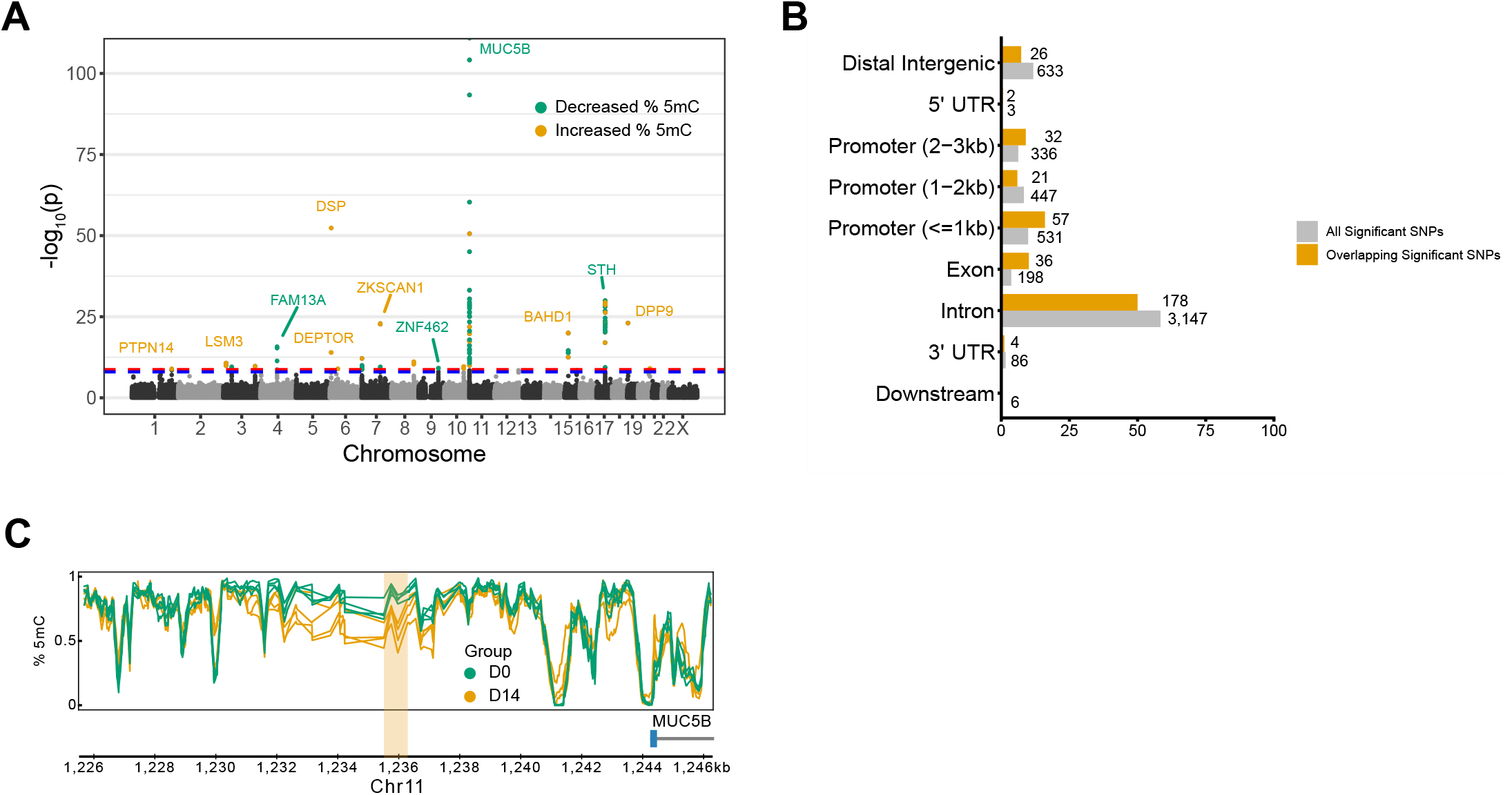
Significant MUC5B GWAS locus overlaps DMR associated with AT2 transdifferentiation. **A** Location of SNPs with direct overlap with a DMR. SNPs associated with DMRs with decreases in methylation are shown in green, and those associated with DMRs with increased methylation are shown in yellow. **B** Distribution of the proximity of SNPs to genomic region annotations. The number of sites for each category is shown next to the respective bar. Annotations for all significant SNPs are shown in gray and annotations for significant SNPs overlapping a DMR are shown in yellow. **C** Intergenic genomic track proximal to *MUC5B*. DMR overlapping a significant disease-associated variant is shown in yellow.

---

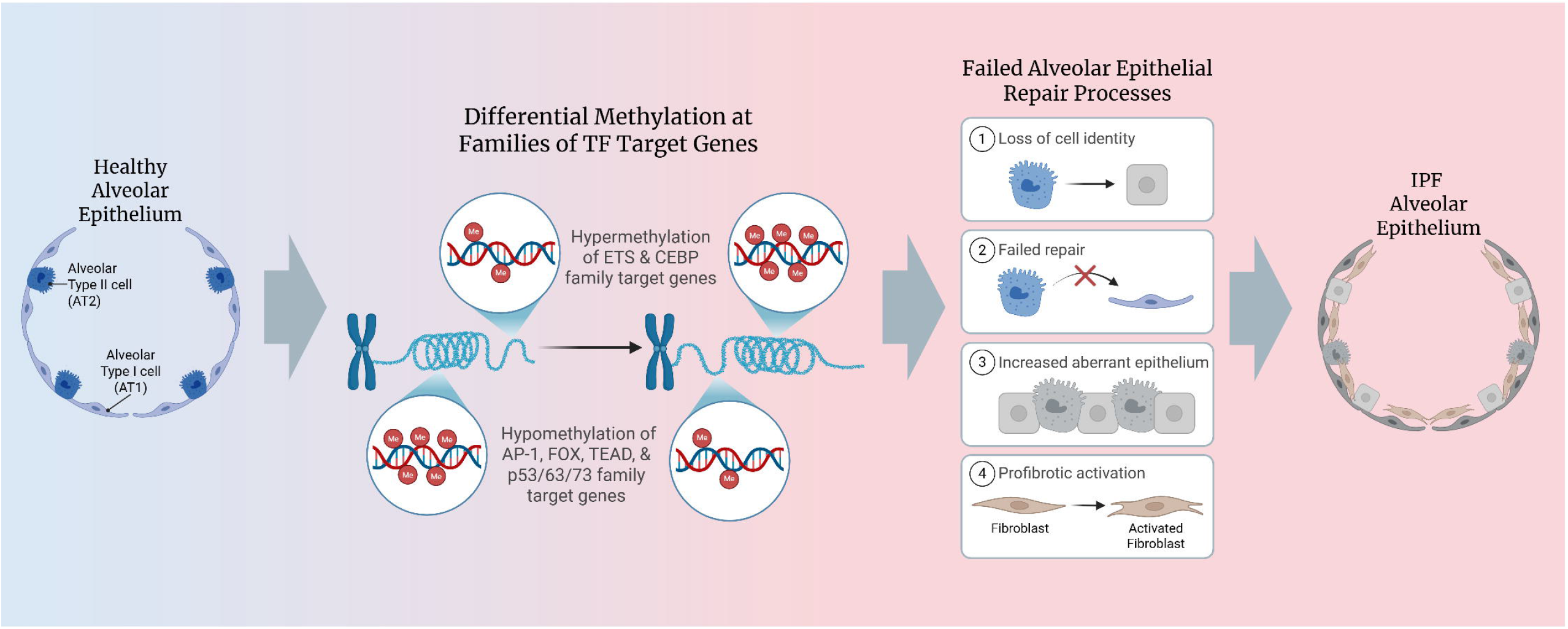

